# Single molecule Grb2 dynamics reveal spatiomechanical modulation of RTK signaling across length scales

**DOI:** 10.64898/2026.10.06.757133

**Authors:** Kashish Jain, Dhruv Thakar, Kshitiz Parihar, Jenny J.Y. Lin, Mary-Kate Hayward, Moses Musiime, Jonathon N. Lakins, Scott D. Hansen, Ravi Radhakrishnan, Jay T. Groves, Valerie M. Weaver, Shalini T. Low-Nam

**Affiliations:** Department of Surgery, University of California, HSE 560, San Francisco, CA 94143; Center for Bioengineering and Tissue Regeneration, University of California, San Francisco, San Francisco, CA, USA; Department of Chemical and Biomolecular Engineering, School of Engineering and Applied Science, University of Pennsylvania, Philadelphia, PA, USA; Department of Chemistry, University of California, 408A Stanley Hall, Berkeley, CA 94720; Department of Chemistry and Biochemistry, University of Oregon, Eugene, OR 97403; Department of Bioengineering, School of Engineering and Applied Science, University of Pennsylvania, Philadelphia, PA, USA; Department of Bioengineering and Therapeutic Sciences, Eli and Edythe Broad Center of Regeneration Medicine and Stem Cell Research, Helen Diller Family Comprehensive Cancer Center, University of California San Francisco, San Francisco, CA 94143, USA; James Tarpo, Jr. and Margaret Tarpo Department of Chemistry, Purdue University, 560 Oval Drive, West Lafayette, IN 47907; Department of Physics, Purdue University, 560 Oval Drive, West Lafayette, IN 47907

## Abstract

Extracellular matrix (ECM)-mediated cell adhesion and tension regulate development and disease through crosstalk with Receptor Tyrosine Kinases (RTKs). RTKs stimulate proliferation, survival, and metabolic reprogramming via Ras/ERK and PI3K/Akt signaling. Emerging evidence identify the biophysical and biochemical integration of integrin-mediated adhesions with RTK signaling, however, the molecular mechanisms regulating this phenotype remain poorly understood. Here, we demonstrate that ECM stiffness potentiates ligand-mediated epidermal growth factor receptor (EGFR) signaling at scales ranging from the single adaptor protein, Grb2 upward to multicellular spheroids and tissues. We show that ECM stiffness-driven RTK signaling is mediated by integrin adhesions that reshape the membrane to induce the formation of RTK signaling hotspots distributed globally across the cell membrane. A parsimonious computational model and biochemical membrane perturbations causally implicate membrane curvature and topography in tension-dependent integrin-EGFR crosstalk. The results demonstrate how the mechanical properties of the ECM direct molecular recruitment and confinement of signaling molecules to regulate growth factor-dependent RTK signaling. Given that many RTK-driven solid tumors develop a stiff stroma, the findings could explain why many therapeutic tyrosine kinase inhibitors develop clinical resistance. The findings suggest clarifying how tension regulates transmembrane receptor signaling could identify unique force-dependent molecular regulators towards which new antitumor therapies could be applied.

## Introduction

Adhesion is the central organizing principle that directs development and maintains tissue homeostasis [1–3]. Cell-cell and cell-extracellular matrix (ECM)-adhesion dictate access to chemical cues and initiate signal integration critical for mediating cell growth, survival, migration, and invasion [1, 4, 5]. Adhesion-directed multicellularity demands that cells continuously respond to biochemical cues within an organized tissue and that they translate inputs detected at the single-molecule level across increasingly longer length scales up to the tissue and organ levels [6–8]. Mechanisms for fidelity in decision-making include macromolecular assembly and formation of specialized signaling domains [9–11]. Thus, proteins spanning and proximal to the membrane and the membrane composition and topography themselves are candidate regulators for signaling outcomes. It has been increasingly clear that information relay can occur over long distances and that local, initiating signals can generate long range effects [12–14]. Receptor tyrosine kinases (RTKs) function within these adhesion-dependent signaling programs, and their activity depends on translating signals that are tuned by the biochemical composition and organization of the ECM [15–17]. However, how adhesion organizes RTK signaling scaffolds remains unclear.

EGFR (erbB1) exemplifies how signaling events are rapidly organized on nanoscale scaffolds. This prototypical RTK is frequently overexpressed, and mis-regulated in many cancers including non-small-cell lung cancer (NSCLC), glioblastoma, pancreatic, and breast cancer [18–20]. Ligand binding induces receptor dimerization and clustering, leading to phosphorylation and recruitment of Grb2–SOS complexes that activate Ras and downstream MAPK/Akt pathways [21–25]. These reactions occur within seconds-to-minutes of EGF binding and generate membrane-associated assemblies. Once activated, individual guanine nucleotide exchange factor (GEF) SOS molecules can processively activate many Ras GTPases and, thus, generate substantial signal amplification [26–29]. An arsenal of anti-EGFR antibodies and small molecule tyrosine inhibitors that target the receptor and many of its downstream molecules including Raf and MAPK has been developed as cancer therapeutics [30–33]. Unfortunately, while these treatments initially show impressive anti-tumor responses, resistance consistently arises [34, 35]. Resistance has been attributed to the emergence and selection of cells expressing mutant EGFR or downstream signaling molecules [36–39]. However, studies have suggested that ECM stiffness and integrin signaling can significantly influence the efficacy of anti-EGFR signaling treatments implicating cell and tissue tension as key modifiers of this pathway [40, 41]. We sought to reconcile how environmental sensing and signaling are intertwined in the integrin-EGFR circuit.

Actomyosin tension links ECM stiffness to adhesion-dependent morphogenesis by coordinating invasive and migratory cell behaviors downstream of activated RTKs. In cancer, ECM stiffening elevates cellular tension to disrupt tissue organization, generating a striking heterogeneity in RTK signaling and driving proliferation, growth, survival, and invasion of individual cells [42–46]. Integrins function as force integrators that couple extracellular mechanics to cytoskeletal and signaling networks, positioning them to participate in RTK signaling crosstalk and activity across scales [6, 47–51]. Consistent with this, RTKs including Eph receptors [52–54] and EGFR [7, 8, 55] exhibit sensitivity to ECM stiffness and actomyosin contractility, although whether effects are direct or mediated through adhesion scaffolds and surface organization is unresolved [48, 52, 56–60]. Thus, force-dependent integrin–ECM adhesion provides a compelling mechanism by which single-cell RTK signaling is tuned and governs multicellular behavior, but how the membrane signaling landscape is organized to facilitate sensitive, rapid responses is unknown.

Here, we show that, at the organismal, tissue, and cellular levels, ECM stiffness modulates EGFR signaling by regulating integrin-dependent cytoskeletal and membrane organization. This framework suggests that the plasma membrane is an active signaling platform that controls activities across molecular and time scales and may be the key source of rapid and sensitive signal integration. We developed a single-molecule Grb2 imaging strategy to quantitatively monitor membrane-proximal signaling immediately downstream of the activated EGFR. Grb2 recruitment to the membrane was used to create a spatially resolved map of receptor activity upstream of SOS recruitment and Ras activation. This strategy enabled localized analysis of the connectivity between ligand-mediated activation of the receptor and recruitment of adaptor proteins leading to Ras activation. This can be considered an impulse-response function for the receptor that maps ligand binding and receptor activation to phosphorylation and priming of downstream signal propagation [61]. We show that EGFR signaling is finely tuned by mechanical properties of the microenvironment at the level of the receptor itself. Remarkably, this modulation can occur over long distances, suggesting global changes in the membrane landscape contribute to integrin-RTK crosstalk and without a requirement for receptors to be colocalized. To explore the contribution of the actin cytoskeleton to long-range signal modulation, we developed a model to map N-WASP-membrane engagement to overall topography and protrusivity for generating topographic sites of specialized activity. Coordinated signaling and membrane organization changes controlled not just signal amplitudes but also durations. While additional layers of signal crosstalk may exist further downstream [7, 8], these data provide an argument for cellular tension altering signaling at the local level of the plasma membrane landscape by physically altering Grb2-EGFR interactions, which, thereafter, affect the global cellular response.

## Results

We interrogated the crosstalk between ECM sensing by cells and mitogenic signaling through the MAPK pathway. Our aim was to elaborate upon the idea that pathways activated by RTKs could be preconditioned by their mechanical and adhesive environment through transmembrane receptors such as integrins [15–17]. We reasoned that if adhesive cues tune RTK activation thresholds and signal propagation, even in the absence of ligand, this would raise fundamental questions about how the mechanical sensing machinery imposes mechanistic changes on receptor behavior at every scale from tissue and organismal levels down to single RTKs. Accordingly, we employed a multiscale view, starting at the tissue scale and moving down to the single molecule level to measure how integrin-mediated sensing imprints directly upon upstream EGFR signaling.

### Tissue-level and multicellular modulation of ECM stiffness affects RTK signaling via Ras

Breast cancer progression is driven by elevated RTK signaling-dependent activation of Ras-stimulated ERK and PI3 kinases that mediate tumor cell growth, survival and invasion [42–46]. The tissue stroma of patients and genetically engineered mouse models (GEMMs) with breast cancer is characterized by a stiffened stroma [7, 8]. Stromal stiffening destabilizes cell-cell adhesions and enhances cell-ECM adhesions to disrupt tissue organization [7, 8, 62, 63]. Experimental breast cancer progression in GEMMs can be reduced by inhibiting the activity of a key collagen crosslinking enzyme lysyl oxidase (LOX) which decreases focal adhesion activity and RTK signaling [7, 8, 62–64]. Given prior work implicating ECM stiffness in EGFR-dependent ERK signaling in cultured organoids, we examined LOX-inhibited PyMT mammary tumors to confirm that reducing stromal stiffness in breast cancer tissues influences integrin-dependent Ras-MAPK signaling (Fig. 1A, B). STIFMaps, a convolutional neural network (CNN)-based machine learning (ML) method that spatially quantifies stromal stiffness in tissues [46], demonstrated that the fibrotic stroma in the PyMT mammary tumors was stiff and that inhibiting LOX-mediated collagen crosslinking prevented stromal stiffening (Fig. 1B,C,E, Fig. S1). Immunofluorescence staining for phospho-FAK revealed that while the mammary epithelium in the control PyMT mice showed robust staining these levels were significantly lower in the LOX-inhibited (BAPN-treated) mammary tissues (p<0.001, unpaired t-test), indicative of reduced integrin-dependent mechanosignaling (Fig. 1D, E). To analyze the effect of reduced tissue stiffness on Ras signaling, we immunostained for downstream Ras effectors phospho-ERK and phospho-AKT substrate [65]. We observed a significant impact with a 34% reduction in phospho-ERK positive nuclei (p<0.05, unpaired t-test) and a 51% decrease in phospho-AKT substrate levels (p<0.001, unpaired t-test; Fig. 1D, E) in the mammary tissues of the LOX inhibitor treated mice, consistent with the hypothesis that a stiffer ECM potentiates Ras signaling.

**Figure 1.**
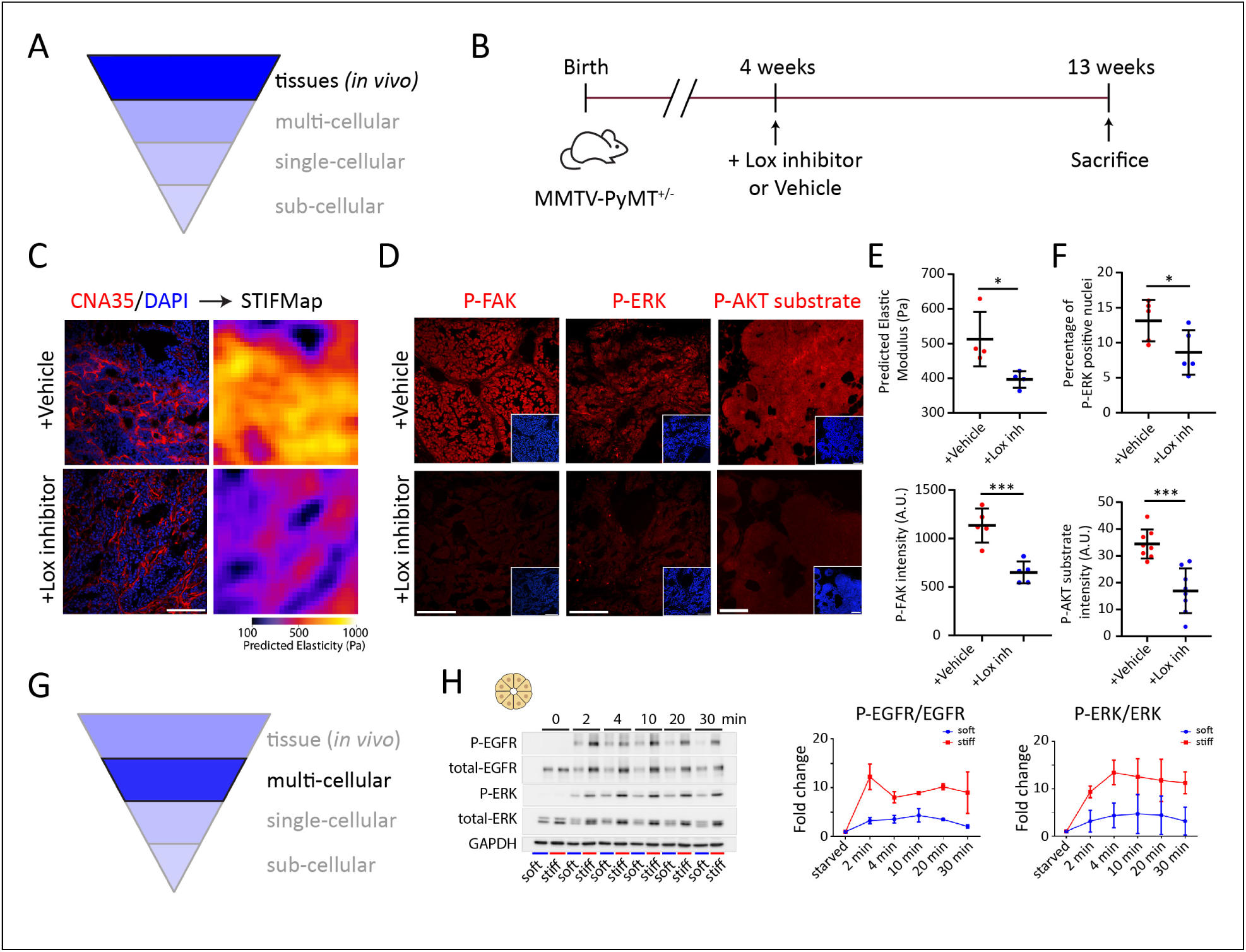
Modulation of environmental stiffness at the tissue and multicellular levels alters RTK signaling. A) Schematic of multiscale signal integration downstream of RTKs, highlighting the tissue scale examined in subsequent panels. B) Experimental design of *in vivo* model. C) (left) Representative immunofluorescence images of vehicle-treated or Lox inhibitor-treated mammary tissues stained for collagen (CNA35, red) and nuclei (DAPI, blue), and (right) the predicted matrix elasticity “collagen paint” (STIFMap, colormap, fire) for the images on the left. Scale bar, 100 μm. D) Representative immunofluorescence for vehicle and Lox inhibitor-treated mammary tissues with antibodies to phospho-FAK (P-FAK), phospho-ERK (P-ERK), and phospho-AKT (P-AKT) substrate (red). Inset shows respective nuclei stained with DAPI (blue). All images are co-scaled. Scale bars, 100 μm. E) Predicted elastic modulus (top) and phospho-FAK (bottom) quantification for images in (C). N = 4 for STIFMap, N = 5 for P-FAK. F) Percentage of phospho-ERK (top) positive nuclei and phospho-AKT (bottom) intensity. N = 5 for P-ERK, N = 8 for P-AKT. G) Schematic highlighting the multi-cellular responses in subsequent panels. H) Immunoblots (left) and corresponding quantification (right) of ratio of P-EGFR:total EGFR, and P-ERK:ERK in EGF-treated MCF10A MECs on soft and stiff Col/rBM gels. N = 3 for both conditions. Data are plotted with biological replicates as scatter points with mean±s.d. Statistical significance was measured by unpaired *t*-test; *p* values: ^∗^*p* < 0.05; ^∗∗^*p* < 0.01; ^∗∗∗^*p* < 0.001; ^∗∗∗∗^*p* < 0.0001; n.s., not significant.

We next used a mammary epithelial cell (MEC) spheroids to map the impact of ECM stiffness on Ras signaling in the context of a multi-cellular model (Fig. 1G, H). Spheroids from the non-malignant MEC line, MCF10A, were embedded within 3D collagen type 1/reconstituted basement membrane (Col/rBM) gels, and the stiffness of the hydrogels was increased from 355±190 Pa (Fig. S1; here, soft) to 2350±1123 Pa (Fig. S1; here, stiff), using non-metabolizable ribose-mediated crosslinking, as confirmed by rheology [43, 66]. Western blot analysis of MEC spheroids embedded in soft (untreated) and stiff (ribose-treated) Col/rBM gels revealed an ∼10-fold increase in EGFR phosphorylation and ERK phosphorylation upon EGF stimulation (Fig. 1H). Consistent with prior studies, elevated ERK phosphorylation was sustained even 30 minutes after stimulation (Fig. 1H) [8]. The increased amplitude and duration confirmed that stiffness-dependent changes in Ras signaling are sustained at the cell population level and raise the question as to how this crosstalk is mediated at the molecular level.

### EGF stimulation leads to long-dwelling membrane recruitment of Grb2

The impact of integrin organization and signaling on RTK signaling suggested that the spatial organization of these transmembrane receptors at the cellular surface mediated the crosstalk between integrin signaling and RTK signaling. Short-range interactions facilitate rapid responses. Therefore, we focused our analysis on early signaling events proximal to the plasma membrane (Fig. 2A). We first validated the EGF-driven RTK signaling at the cellular and subcellular scales using membrane-selective imaging in a Total Internal Reflection Fluorescence (TIRF) configuration (Fig. 2A-C). Growth factor binding to receptor tyrosine kinases (RTKs) leads to their phosphorylation, promoting to the recruitment of adaptor proteins such as Grb2 and SOS, which activate Ras, leading to the activation of downstream kinases such as ERK, MAPK, AKT and their associated signaling networks (Fig. 2B) [16, 67]. Tyrosine phosphorylation at the 1068 residue acts as a primary site for Grb2 binding to EGFR, with tyrosine phosphorylation at the 1086 residue acting as a secondary binding site [68, 69]. Grb2 recruitment to phospho-tyrosine residues on the cytoplasmic tail of EGFR is, thus, the most proximal readout of receptor activation (Fig. 2B) [67, 70]. Receptor phosphorylation may be primed by ligands or microenvironmental inputs [43, 71, 72]. We employed single molecule detection of Grb2 docking as an early readout of the EGFR impulse-response function that proceeds through MAPK signaling via Ras to regulate cell growth, survival, and invasion and migration [65, 67].

**Figure 2.**
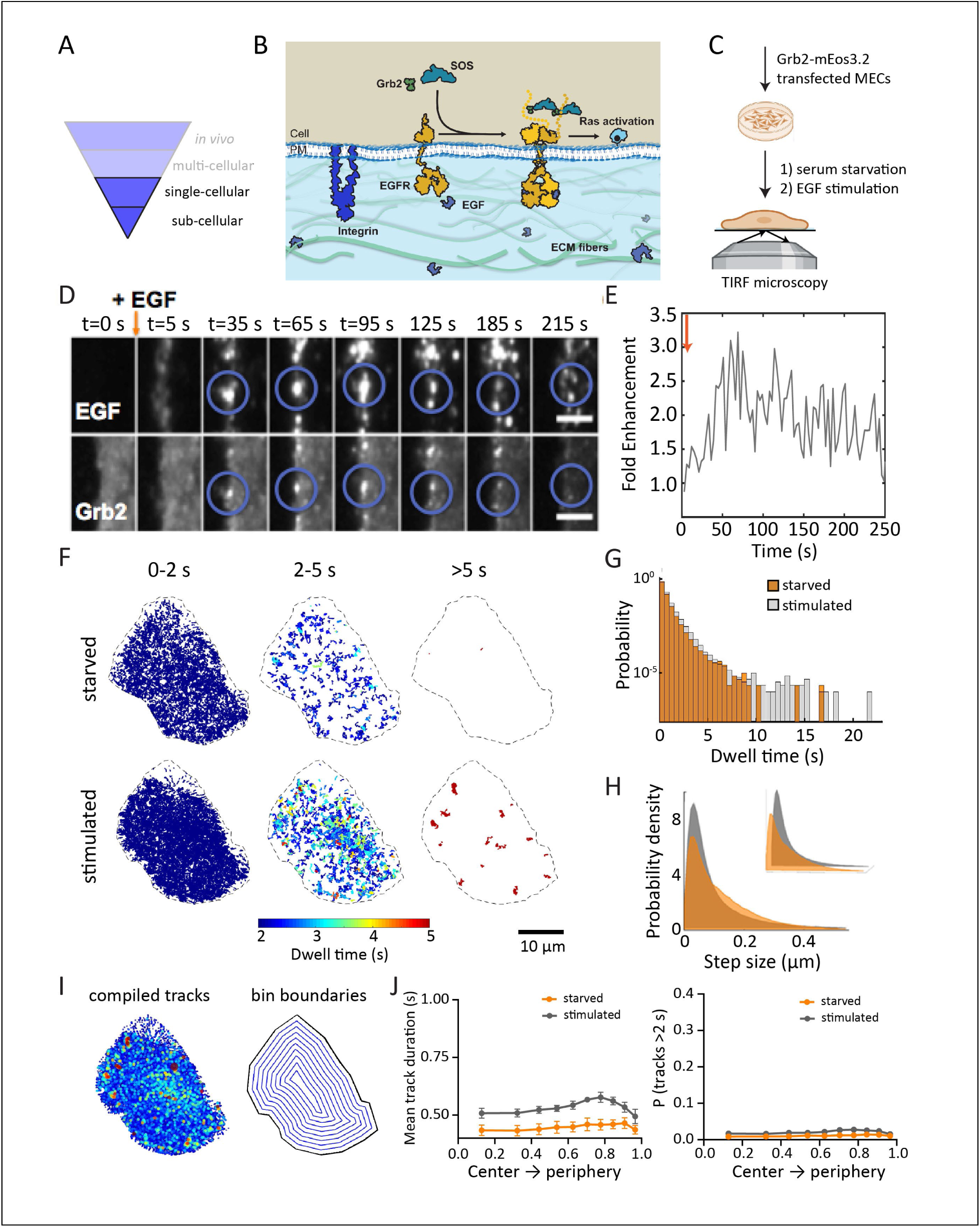
Grb2 recruitment is a membrane-proximal, early activation readout for EGF-dependent RTK signaling. A), Subsequent panels refer to single cell responses. B), Schematic of MAP kinase signaling cascade downstream of EGF-RTK interaction. EGFR activation by EGF binding induces a conformational change, leading to its dimerization and tyrosine phosphorylation, recruiting Grb2. Grb2 binding recruits SOS, wherein it becomes catalytically active and activates Ras. Propagation of MAPK signaling downstream of Ras leads to ERK activation. C) Schematic of MEC transfection, seeding, stimulation, and TIRF imaging. D) Representative montage of mEos3.2-Grb2-expressing MEC cell stimulated with 100 ng/mL fluorescent EGF. The blue circle indicates a strong, rapid recruitment of Grb2 at sites where EGF is bound. Scale bar, 5 µm. E) Fold enhancement in Grb2 recruitment following EGF stimulation. In (D) and (E), the orange arrow indicates the time of EGF addition. F) Ensemble projection of dwell times for mEos3.2-Grb2 at the PM on fibronectin-coated glass substrates under starved or EGF-stimulated conditions. Dwells are color-coded based on duration and plotted over short, moderate, and long-dwelling regimes. Scale bar, 5 µm. G), Dwell time distributions for mEos3.2-Grb2 on fibronectin-coated glass substrates. H) Step size distributions for mEos3.2-Grb2 mobility shown enlarged in 2D with an inset showing the same distributions in 3D. In (G) and (H), the colored distribution represents dwell times under starvation conditions, and the gray distribution indicates dwell times following stimulation with EGF. N>5 cells across 5 experiments. I) Spatial analysis for the Grb2 dwell times for cells on glass. All Grb2 dwell events were superimposed on the cell boundary, with each event represented as a circle whose size and color reflected dwell time. The cell was divided into 10 concentric bins, and the bin boundaries shown. J) The mean track duration and probability of long events was plotted as a function of the bins, from the cell center to cell periphery. N = 5 cells for each condition.

To examine how growth factor treatment affected Grb2 recruitment to the membrane at the molecular level, we treated MCF10A cells, engineered to express a mEos3.2-labeled Grb2, treated with fluorescently labeled EGF after serum starvation (Fig. 2C). MECs transiently expressing Grb2-mEos3.2 were plated on glass substrates coated with fibronectin, an ECM ligand overexpressed and deposited in the stroma of the majority of solid tumors [73–76], serum starved and imaged using TIRF microscopy. We observed an increase in Grb2 fluorescence intensity at the membrane, which overlapped with fluorescent EGF (Fig. 2D). This increase in Grb2 colocalization with ligated receptors was virtually instantaneous (Fig. 2D), indicating a rapid RTK activation. We quantified the enhancement of Grb2 with respect to EGF and observed a ∼3-fold enhancement of Grb2 within a minute of EGF stimulation (Fig. 2E). The clustered EGF signal likely reflected ligand binding, receptor dimerization and tyrosine phosphorylation at residues 1068 and 1086, and some EGFR oligomerization [16, 67]. These phosphorylated sites primed Grb2 binding, thus validating the use of Grb2 recruitment as a readout of RTK signaling.

We further characterized individual Grb2 recruitment and dynamics at the cell membrane using single particle tracking photoactivated localization microscopy (SPT-PALM) [11, 77, 78]. The dynamics of single, membrane-localized mEos3.2-Grb2 molecules were measured using short exposures of 405 nm excitation, which stochastically converted a subpopulation of mEos3.2-Grb2 molecules to the 561 nm excited state (Fig. S2), together with high-speed imaging. Cycles of photoconversion and single molecule imaging enabled sampling of Grb2 recruitment dynamics for thousands of molecules over many minutes. SPT-PALM analysis of the MECs revealed that Grb2 molecules frequently arrive across the membrane for short durations of up to 2 seconds, even in the absence of ligand (Fig. 2F, G). Molecular binding dwell times exceeding 5 seconds were rare, suggesting a basal signal amplification occurred without the need for longer docking times. Just as with the visualization of EGF binding, ligand stimulation led to an increased number of Grb2 docking events at the membrane and a marked increase in long dwell times, including events with durations over 5 sec (Fig. 2F, G). Tracking of these long dwelling events demonstrated that long-dwelling Grb2 was restricted to small regions all across the cell surface (Fig. 2F). These areas are consistent with the assembly of signaling hotspots which could lead to prolonged Ras activation [11, 77–81]. We further quantified the lateral diffusion of Grb2 molecules after their recruitment to the membrane (Fig. 2H). The diffusion step-size histogram for the stimulated condition shifted to the left compared to the starved condition (Fig. 2H), indicating that RTK activation led to shorter Grb2 steps and a slowed diffusion.

Since we observed long dwelling Grb2 events restricted to specific regions across the cell membrane, we further performed analysis of the spatial organization of these tracks. Dwells of duration exceeding 3 seconds were first transformed to their center-of-mass (Fig. 2I). A binary cell mask was then used to divide the cell into 10 concentric bins of equal radius, from the center to the periphery (Fig. 2I), and the Euclidean distance of each long dwell was calculated from the edge and mapped to the bins. Simulations for a heterogeneous distribution with longer events at the periphery exhibited a progressive increase in probability of long dwells based on bin distance from the center of the contact (Fig. S3), whereas simulations for a homogeneous distribution showed a relatively constant probability across the radial bins (Fig. S4). We observed that the mean track duration was higher for stimulated cells as compared to the starved cells, but that neither condition showed an appreciable location-dependent mean dwell time (Fig. 2J). These data suggest that there was a homogeneous distribution of long dwelling Grb2 events across the cell surface.

### Fibronectin concentration-regulated integrin signaling affects long Grb2 dwelling events

Since Grb2 could be used as a robust readout of RTK signaling (Fig. 2), we next investigated the effect of modulating integrin signaling on RTK signaling at the membrane using our Grb2 SPT-PALM strategy. Previous evidence of the crosstalk between the adhesome components and RTK signaling indicates that integrin signaling could regulate EGFR signaling at the cellular level [82–84]. However, a conclusive mechanism by which this crosstalk occurs has remained elusive. Therefore, we systematically tested the role of integrin adhesion formation on RTK signaling. We first controlled the extent of integrin signaling by changing the concentration of the extracellular ligand, fibronectin, adsorbed on glass substrates before MEC seeding (Fig. S5.). Similar to previous studies [85, 86], we observed that increasing fibronectin concentration increased an integrin-dependent cell phenotype that included a 5-fold increase in cell spreading area when fibronectin concentration was varied over 2 orders of magnitude (Fig. 3A, B). We further stained for activated focal adhesion kinase (FAK phosphorylated at tyrosine 397; pFAK), to measure effects on integrin-dependent signaling. We observed an increase in pFAK adhesion area as a function of fibronectin concentration (Fig. 3A, C, Fig. S6A), demonstrating how increasing the fibronectin concentration directly enhanced integrin signaling in MEC cells. The adhesions formed primarily close to the cell edge, similar to prior demonstrations [85, 86].

**Figure 3.**
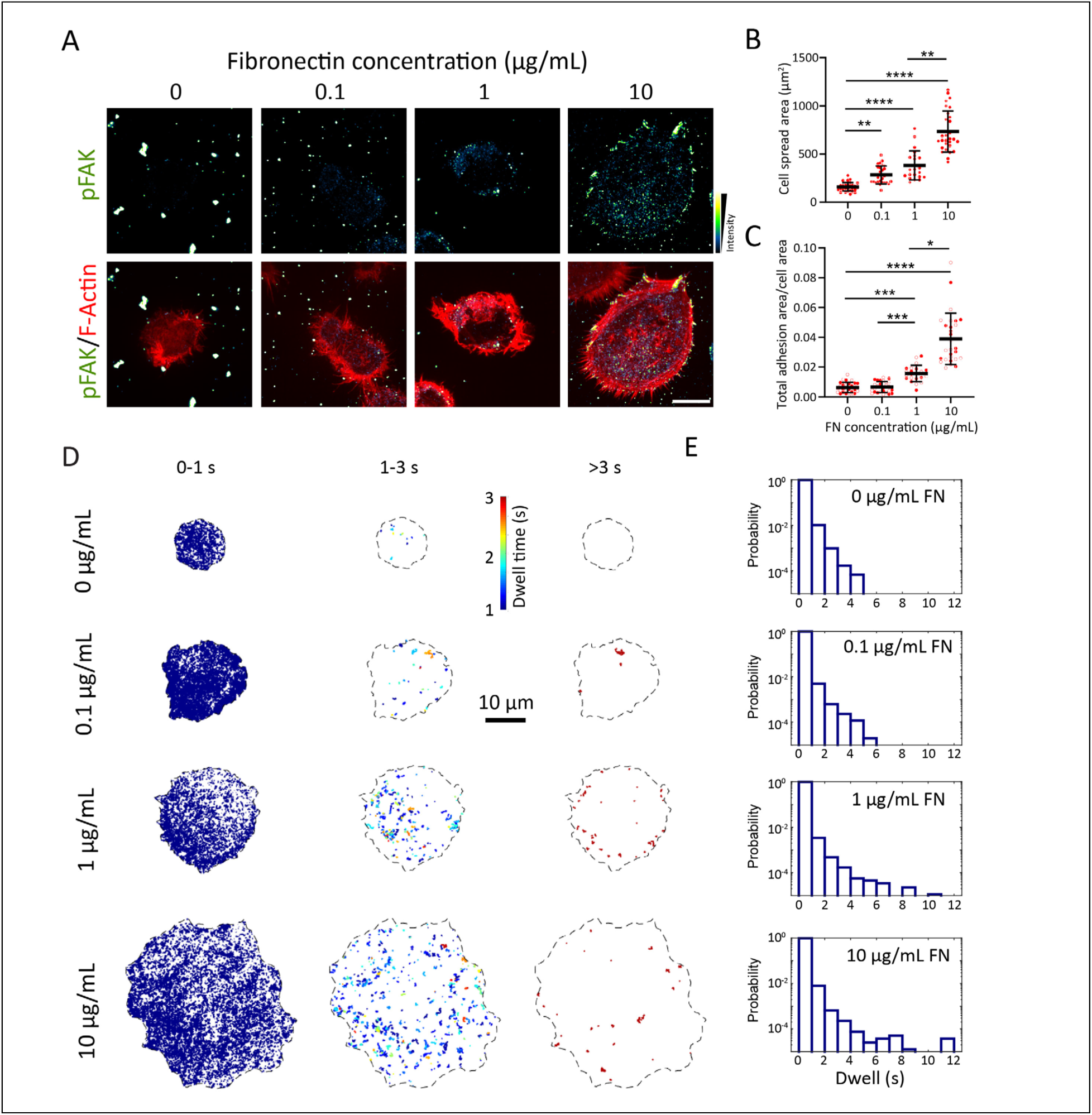
Fibronectin concentration-mediated integrin signaling affects RTK signaling. **A)** MCF10A cells plated on glass substrates coated with different concentrations (0, 0.1, 1, or 10 µg/mL) of fibronectin and allowed to adhere for 4 hours (under serum starvation conditions). Immunofluorescence micrographs of Focal Adhesion Kinase (FAK)-phospho-Y397 (pFAK) to visualize adhesions, and a merge of actin and pFAK. Scale bar, 10 µm. **B**) Plot of cell spread area as a function of fibronectin concentration and **C**) total pFAK area normalized to cell spread area. Data is represented as mean ± standard deviation along with all individual values (n ≥ 25 cells from three independent experiments, analyzed by the Kruskal-Wallis one-way ANOVA and Dunn’s post hoc test; \**P*<0.05, \*\**P*<0.01, \*\*\**P*<0.001, \*\*\*\**P*<0.0001). **D)** Representative spatial mapping of the mean mEos3.2-Grb2 dwells on different concentrations of fibronectin-coated glass substrates. Dwells are color-coded based on duration and plotted over short (0-1s), moderate (1-3s), and long-dwelling (>3s) regimes. Scale bar, 10 µm. **E)** Dwell time probability distributions for single molecule mEos3.2-Grb2 recruitment to the membrane as a function of fibronectin concentration on glass. All distribution represents dwell times following stimulation with EGF. N > 5 cells across 3 experiments.

We next performed SPT-PALM on Grb2-mEos3.2 and quantified the dwell time of Grb2 recruitment events at the membrane. The majority of Grb2 tracks had a dwell time of less than 200 ms, and Grb2 binding events of longer duration (>3 sec) were observed at the higher fibronectin densities 1 µg/mL and 10 µg/mL (Fig. 3D, E). These longer dwelling events were spatially confined, suggesting the formation of signaling clusters, where robust, prolonged signaling takes place. Since these long dwelling events were observed only in high fibronectin concentrations, this suggests an effect of integrin signaling on Grb2 membrane recruitment and cluster formation above a critical threshold.

### Grb2 recruitment to the cell surface is enhanced on stiff substrates

To further elucidate the crosstalk between integrin and RTK signaling, we next modulated substrate stiffness as an independent means to alter integrin signaling at constant densities of fibronectin. We chose polysiloxane gels, which are compatible with TIRF imaging because their refraction index is close to glass, which allows total internal reflection at the gel-cell interface, so that events at the ventral membrane could be visualized with single molecule resolution [87–89]. The ratio of polymer to crosslinker was adjusted to control stiffness by a factor of approximately one order of magnitude. The elastic moduli, characterized by shear rheology, covered two extremes: ∼250 Pa (soft) and ∼10 kPa (stiff) [7, 88]. This spectrum covers environmental stiffnesses characteristic of normal and transformed breast tissue, respectively [7]. The gels were spin-coated onto glass coverslips, the gel surface was silanized, and 1 μg/ml of fibronectin was crosslinked to the surface (Fig. S6.) [87–89]. Similar to prior reports, MCF10A MECs on soft gels exhibited smaller cell spreading areas and aspect ratios close to 1 (Fig. 4 A-C, Fig. S7). In contrast, cells on stiff substrates were more elongated with a 3.5x higher cell spreading, indicating a morphological response to stiffness tuning (Fig. 4 A-C, Fig. S6B) [8, 88]. We also observed a 78% increase in adhesion area and adhesion length on the stiff substrates compared to the soft substrates (Fig. 4 A-C, Fig. S6B).

**Figure 4.**
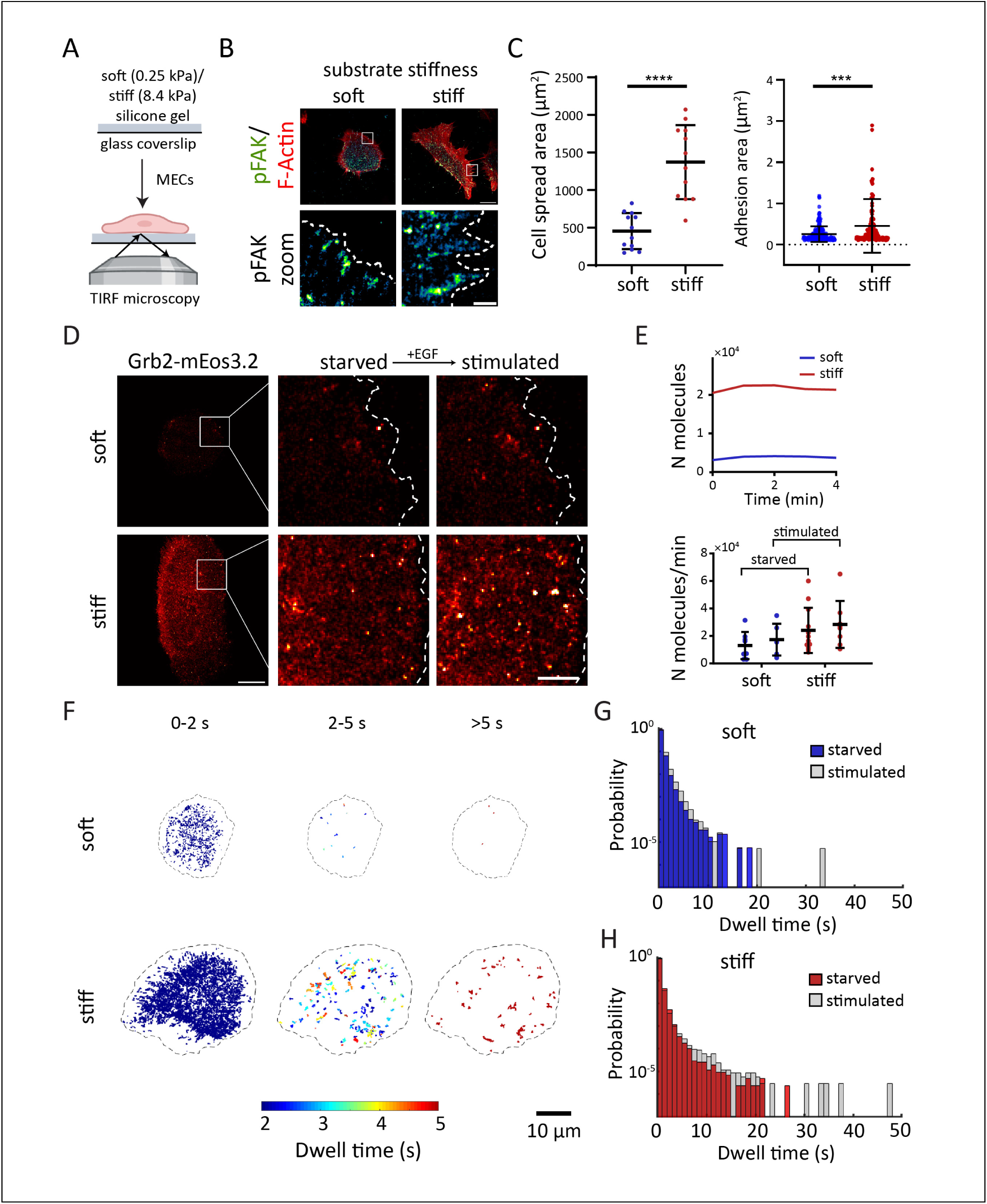
Substrate stiffness-mediated integrin signaling affects RTK signaling. **A)** Schematic of MEC seeding on soft and stiff silicone gels and TIRF imaging. **B**) MCF10A cells plated on soft and stiff silicone gels coated with fibronectin and allowed to adhere for 4 hours. Immunofluorescence micrographs of Focal Adhesion Kinase (FAK)-phospho-Y397 (pFAK) to visualize adhesions, and a merge of actin and pFAK, along with a zoom of the white box. Scale bar (top), 10 µm. Scale bar (bottom), 2 µm. **C)** Plot of cell spread area and pFAK adhesion area for cells spread on soft vs stiff gels. Data is represented as mean ± standard deviation along with all individual values (n ≥ 10 cells from three independent experiments, analyzed by unpaired *t*-test; *p* values: \*\*\**P*<0.001, \*\*\*\**P*<0.0001). **D)** Super-resolution rendered images of MECs on soft vs stiff silicone gel. Region of interest enlarged following 4 hours of serum starvation (Starved) and subsequent 4 minutes of stimulation with 100 ng/mL EGF (Stimulated). Scale bar, 10 μm, 2 μm (zoom). **E)** Quantification of the number of Grb2 molecules recruited during the time course of stimulation (top), and. Single cell distributions of number of Grb2 molecules recruited per minute under starvation or stimulation conditions (bottom). N > 5 cells each. **F)** Representative spatial mapping of the mean mEos3.2-Grb2 dwells on soft vs stiff substrates. Dwells are color-coded based on duration and plotted over short (0-2s), moderate (2-5s), and long-dwelling (>5s) regimes. Scale bar, 10 µm. **G, H)** Dwell time probability distributions for single molecule mEos3.2-Grb2 recruitment to the membrane on a soft (G) and a stiff (H) substrate. The colored distribution represents dwell times under starvation conditions, and the gray distribution indicates dwell times following stimulation with EGF. N > 5 cells, individually matched before and after EGFR, across 3 experiments.

**Fig. 5.**
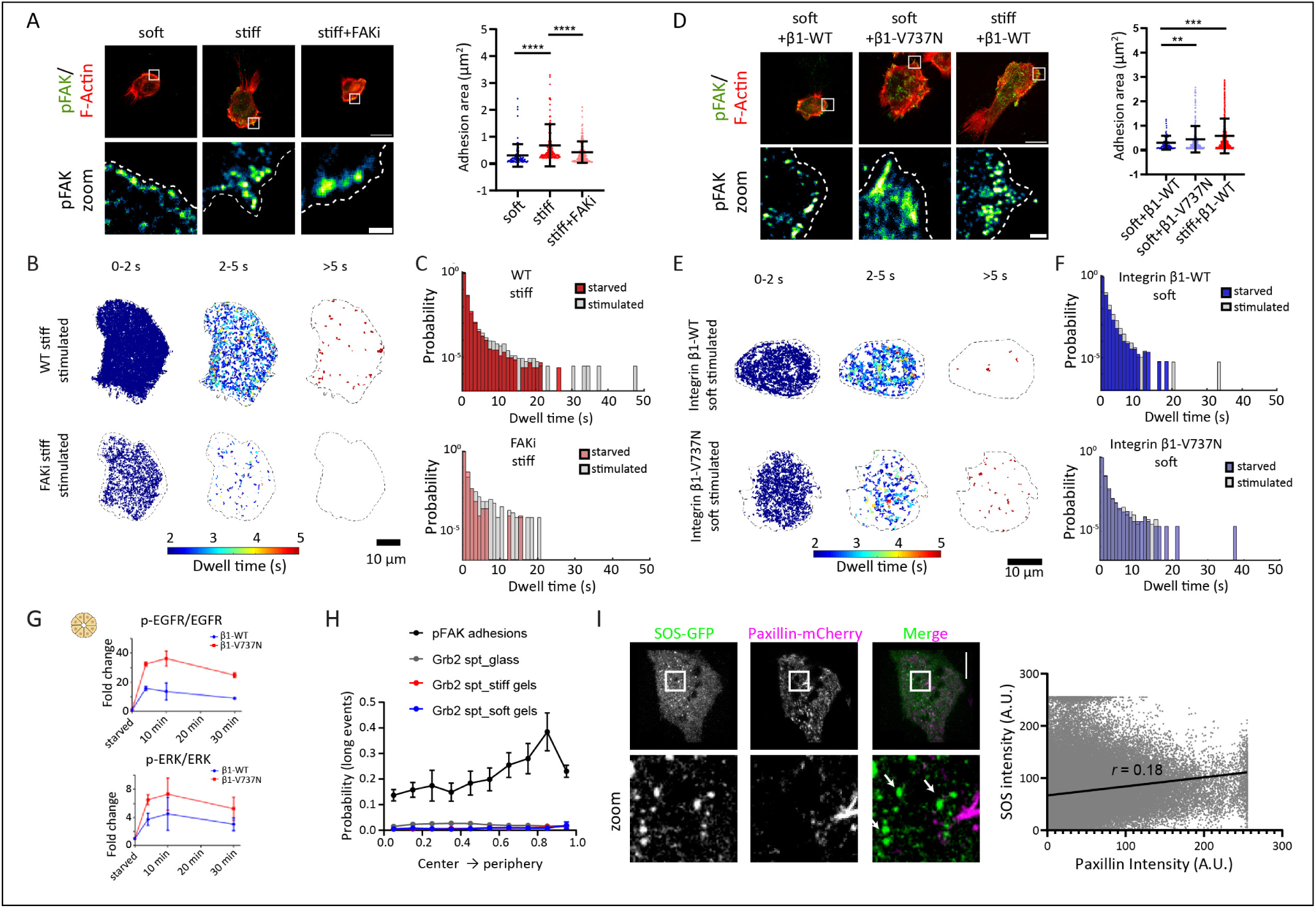
Alterations in integrin organization and signaling potently modulate RTK activation, but not by direct interaction. A) Representative images of MCF10A cells plated on soft and stiff silicone gels with or without FAK inhibitor 14 (FAKi), along with quantification of cell spread area and adhesion area. Immunofluorescence micrographs of Focal Adhesion Kinase (FAK)-phospho-Y397 (pFAK) to visualize adhesions, and a merge of actin and pFAK, along with a zoom of the white box. Scale bar (top), 10 µm. Scale bar (bottom), 2 µm. Plot of cell spread area and pFAK adhesion area for cells spread on soft vs stiff gels. Data is represented as mean ± standard deviation along with all individual values (n ≥ 10 cells from three independent experiments, analyzed by unpaired *t*-test; *p* values: \*\**P*<0.01, \*\*\**P*<0.001, \*\*\*\**P*<0.0001). B) Representative spatial mapping of all mEos3.2-Grb2 dwells on stiff substrates with or without the FAKi. Dwells are color-coded based on duration and plotted over short (0-2s), moderate (2-5s), and long-dwelling (>5s) regimes. Scale bar, 10 µm. C) Dwell time probability distributions for single molecule Grb2 recruitment to the membrane on a stiff substrate with or without the FAKi. The colored distribution represents dwell times under starvation conditions, and the gray distribution indicates dwell times following stimulation. N > 5 cells across 3 experiments. D) Representative images of MCF10A cells plated on soft and stiff silicone gels expressing either integrin β1-WT or the V737N mutant, along with quantification of cell spread area and adhesion area. Immunofluorescence micrographs of a merge of actin and pFAK, along with a zoom of the white box. Scale bar (top), 10 µm. Scale bar (bottom), 2 µm. Plot of cell spread area and pFAK adhesion area for cells as indicated. Data is represented as mean ± standard deviation along with all individual values (n ≥ 10 cells from three independent experiments, analyzed by unpaired *t*-test; *p* values: \*\**P*<0.01, \*\*\**P*<0.001, \*\*\*\**P*<0.0001). E) Representative spatial mapping of all mEos3.2-Grb2 dwells on stiff substrates expressing either integrin β1-WT or the V737N mutant. Dwells are color-coded based on duration and plotted over short (0-2s), moderate (2-5s), and long-dwelling (>5s) regimes. Scale bar, 10 µm. F) Dwell time probability distributions for single molecule Grb2 recruitment to the membrane on a soft substrate expressing either integrin β1-WT or the V737N mutant. The colored distribution represents dwell times under starvation conditions, and the gray distribution indicates dwell times following stimulation. N > 5 cells across 3 experiments. G) Quantification of immunoblots for ratio of P-EGFR: total EGFR and P-ERK: total ERK in EGF-treated MECs on soft Col/rBM gels expressing either integrin β1-WT or the V737N mutant.

By combining the TIRF-compatible gels and single-molecule Grb2-mEos3.2 imaging (Fig. 4a), we next analyzed the spatiotemporal features of Grb2 membrane recruitment as a function of extracellular matrix stiffness. We acquired single-molecule recruitment sequences at a rate of 20 Hz for up to 4 minutes on serum-starved MECs. The same cells were imaged for up to 4 minutes immediately following stimulation with saturating amounts of EGF (100 ng/mL). We focused on early time points following EGF exposure to minimize contributions from endocytosis, which has been shown to contribute to EGF-induced signaling [15, 90].

For all cells, stimulation with EGF led to an increase in the number of Grb2 molecules recruited to the surface, indicating that a substantial fraction of Grb2-related behaviors observed reflect the activity of the EGF receptor (Fig. 4 D, E). Recruitment occurred across the adherent surface and was approximately 2-fold higher on stiff substrates compared to the soft substrates (Fig. 4 D, E). However, the rate of Grb2 recruitment did not dramatically increase (Fig. 4E), indicating that another feature of the Grb2 dynamics likely accounts for the differences in signal amplitude. However, total EGFR surface expression, measured in cells following overnight culture, confirmed by flow cytometry, demonstrated a minimal change in membrane receptor expression in the cells on soft or stiff gels (Fig. S7).

### Substrate stiffness affects long Grb2 dwelling events

Since we failed to observe a significant difference in the recruitment of Grb2 to the membrane upon EGF stimulation, we hypothesized that the dynamics, and, not the amount, of Grb2 membrane recruitment events on stiffer substrates could be particularly consequential for regulating MAPK signaling. Furthermore, evidence for Ras nanoclusters on the cell membrane would indicate that there are distinct regions on the membrane that contribute disproportionately to Ras activation [91, 92]. We therefore performed SPT-PALM on Grb2-mEos3.2 and quantified the molecular dwell time of Grb2 recruitment events at the membrane. On stiff gels, the tail of Grb2 dwell times was extended up to tens of seconds following ligand stimulation (Fig. 4 F-H), while on soft gels, there was a negligible increase in dwell times upon EGF exposure (Fig. 4 F-H). Longer events were rare for cells on soft gels, consistent with lesser signal propagation (Fig. 4 F, G). The diffusion step-size histogram for the stiff cells also shifted to the left by a larger magnitude compared to the soft gels, indicating a slower diffusion (Fig. S8). Specificity was verified using a membrane-tethered mEos3.2 construct whose mobility remained unchanged following EGF addition (Fig. S9). We also observed that for cells on both soft and stiff gels, Grb2 membrane dwelling events were distributed across the adherent surface (Fig. 4F, Example of starved cells shown in Fig. S10). Thus, similar to integrin adhesion modulation by changing fibronectin concentration, increasing integrin adhesion formation by increasing substrate stiffness led to an increased Grb2 membrane dwelling, which then resulted in increased RTK signaling.

### Integrin adhesion modulation on soft and stiff substrates affects Grb2 recruitment

To clarify whether the stiffness-dependent effects on RTK signaling were mediated via integrin adhesions, we modulated integrin-dependent focal adhesion assembly and FAK signaling in the cells plated directly on soft and stiff substrates using pharmacological treatments and investigated their effect on Grb2 membrane recruitment and single-particle dynamics. We first treated MECs with an FAK inhibitor (60 nM FAK14; FAKi) to decrease integrin adhesion formation and FAK signaling on stiff substrates (Fig. 5A). We observed a 54% decrease in cell spread area and a 41% decrease in adhesion area with FAKi treatment on stiff substrates, compared to untreated cells (Fig. 5A, Fig. S11A). SPT-PALM analysis demonstrated a decrease in long Grb2 membrane dwell times (Fig. 5B, C) in FAKi treated cells on stiff substrates, resulting in comparable Grb2 dwell times on the soft substrates (Fig. 5B,C, Fig. 4). Thus, decreasing integrin adhesion formation and FAK signaling on stiff substrates compromised RTK signal integration on stiff substrates.

We next investigated if a predisposition towards integrin clustering favored increased RTK signaling on soft substrates. The integrin β1-V737N mutant has been shown to foster focal adhesion assembly and increase adhesion size and FAK signaling in cells plated on soft substrates by enhancing integrin clustering [8]. We observed that MECs expressing the β1-V737N mutant had a 65% higher cell spreading, assembled 46% larger focal adhesions and had elevated phospho-FAK when plated on the soft gels as compared to MECs expressing integrin β1-WT (Fig. 5D, Fig. S11B). We next performed SPT-PALM on MECs expressing either integrin β1-WT or β1-V737N spread on soft gels. We observed that the MECs expressing the integrin β1-V737N had long Grb2 membrane dwell times even when plated on soft gels (Fig. 5E, F), similar in phenotype to MECs seeded on the stiff substrates (Fig. 4). Thus, increasing integrin adhesions and signaling in cells plated on soft substrates led to an increase in Grb2 membrane dwelling, which in turn can facilitate prolonged RTK signaling. To test the effect of the integrin V737N mutation on EGFR activation and downstream ERK signaling, we used the WT vs V737N mutation transfected MCF10A acini embedded in soft gels (Fig. 5G). Western blot analysis demonstrated a 2x higher fold increase in phospho-EGFR normalized to total EGFR in the β1 integrin-V737N expressing acini compared to the β1 integrin-WT expressing acini (Fig. 5G). We observed a similar 2x fold increase in phospho-ERK normalized to total ERK with the β1 integrin-V737N mutant compared to the β1 integrin-WT cells (Fig. 5G). These findings further show that modulating focal adhesions and signaling can regulate stiffness-dependent RTK signaling.

### Enhanced Grb2 dwell does not require close proximity to sites of adhesion

Since RTK signaling was a direct consequence of integrin adhesion formation and adhesion signaling, we investigated the spatial correlation between integrin-associated complexes and RTK signaling complexes. A high spatial correlation would suggest direct interaction between the adhesions and RTK signaling complex, which could provide mechanistic insight. We compared the spatial analysis for Grb2 SPT-PALM with a known heterogeneously distributed pFAK adhesion spatial analysis (Fig. 3A) [85, 86]. For pFAK adhesions, we observed a progressive increase in probability of long events (here, large adhesions > 0.5 µm^2^) from cell center to the periphery (Fig. 5H). This is consistent with previous work which show that mature focal adhesions primarily appear near the cell periphery [85, 86]. We next compared Grb2 SPT spatial analysis for glass, soft and stiff gels with the pFAK spatial analysis. However, and surprisingly, spatial analysis for all Grb2 SPT substrates exhibited a relatively uniform probability across the radial bins, indicating a diminished spatial gradient, and a more homogeneous distribution (Fig. 5H). To further validate the spatial analysis, we co-expressed Paxillin-mCherry and SOS-EGFP in MEC cells and imaged the cells on fibronectin-coated glass substrates. We observed that a majority of SOS clusters failed to colocalize with Paxillin (Fig. 5I), confirming that there is no direct spatial correlation between the two complexes (R^2^=0.03; Fig. 5I). The spatial analysis and correlation analysis showed that there was minimal to no direct interaction between adhesions and the RTK complex and instead suggested that the effect of adhesions on Grb2 membrane dwelling were likely due to a global mechanism.

### Membrane topology is homogeneous and is dependent on integrin adhesions

Membrane receptor dynamics and signal strength have been shown to be regulated by membrane curvature [81, 93, 94], a physical feature of the cell membrane. Thus, we hypothesized that the organization of the membrane could itself modulate Grb2 membrane dwelling and, consequently, RTK signaling outcomes. Membrane curvature has also been shown to be dependent on ECM organization [95], emphasizing the possibility that stiffness-dependent Grb2 dwelling might be membrane curvature-mediated. To explore this possibility, we performed scanning angle interference microscopy (SAIM) and analyzed the spatial probability of membrane curves obtained (Fig. 6A). SAIM utilizes the interference between incident laser and laser reflected from a silicon wafer at the fluorescently labelled cell membrane, which allows for nanometer scale resolution of membrane topology close to the adherent surface (Fig. 6A) [81, 96]. Membrane height maps obtained from SAIM images showed ridges (negative curvature) and grooves (positive curvature) on the membrane in the range of ∼50 nm (Fig. 6A), and spatial analysis of these ridges and grooves demonstrated that they were homogeneously distributed across the adherent membrane (uniform probability across the radial bins; Fig. 6B).

**Figure 6.**
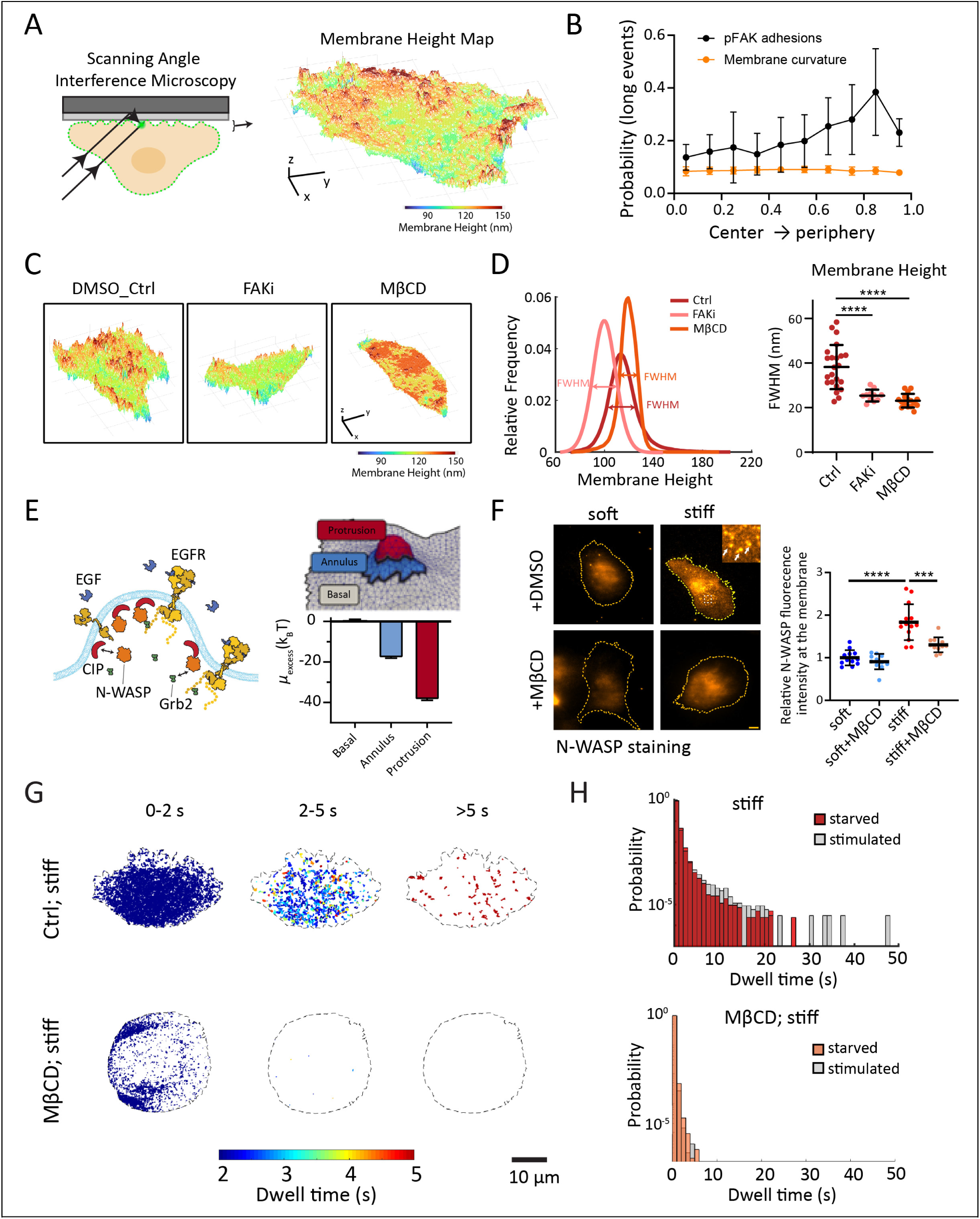
Heterogeneity in RTK activation does not require spatial colocalization with integrins and is impacted by membrane topography. A) Schematic for scanning angle interference microscopy (SAIM) imaging of the cell membrane, along with a 3D rendered image of the ventral membrane. Scale bar, 50 µm in x,y and z axes. B) Plot for spatial analysis with probability of long events (large-scale ridges-groove topography) for membrane curvature and pFAK adhesions as a function of rois from cell center to cell periphery. N=5 cells for each condition. C) Representative 3D rendered ventral membrane height images for MECs treated with DMSO (control), FAKi or with MβCD. Scale bar, 50 µm in x,y and z axes. D) Histogram of the membrane heights for cells shown in C), along with the full width half maxima (FWHM), and the quantification of FWHM for multiple cells on the right. (n > 10 cells from three independent experiments, analyzed by unpaired *t*-test; *p* value: \*\*\*\**P*<0.0001). E) Schematic of curvature-inducing protein (CIP) mediated interaction of N-WASP and Grb2. N-WASP is recruited to the plasma membrane by CIPs, which can then interact with Grb2. (right) Diagram showing spatial regions associated with membrane protrusions including the basal, annulus and protrusion domains on top, with bar graph of free energy, quantified by the excess chemical potential (µ_excess_(k_B_T)) for recruitment of curvature-inducing proteins to the basal, annulus, and protrusion regions on bottom. F) Representative images of MCF10A cells plated on soft and stiff silicone gels and immunostained for N-WASP, imaged using TIRF, along with the intensity quantification. Scale bar, 10 µm. Data is represented as mean ± s.d. along with all individual values (n ≥ 10 cells from three independent experiments, analyzed by one-way ANOVE followed by Tukey’s multiple comparisons test; *p* values: \*\*\**P*<0.001, \*\*\*\**P*<0.0001). G) Representative spatial mapping of all mEos3.2-Grb2 dwells on stiff substrates treated with either DMSO control or with MβCD. Dwells are color-coded based on duration and plotted over short (0-2s), moderate (2-5s), and long-dwelling (>5s) regimes. Scale bar, 10 µm. H) Dwell time probability distributions for single molecule Grb2 recruitment to the membrane on a soft substrate treated with either DMSO or MβCD. The colored distribution represents dwell times under starvation conditions, and the gray distribution indicates dwell times following stimulation. N > 5 cells across 3 experiments.

We further analyzed the effect of integrin adhesion formation on membrane topology, quantified by SAIM. MECs treated with FAKi had a flatter adherent membrane, with a lower number of grooves and ridges, along with a lower intensity (Fig. 6C). We quantified these effects by calculating the Full Width Half Maxima (FWHM) of the membrane height histogram (Fig. 6D) and observed a 34% decrease in the FWHM of membrane height upon FAKi treatment (P<0.0001; Fig. 6D). This reduction in membrane topology when integrin adhesions were reduced suggests that membrane topology is controlled by integrin adhesions, which could be a homogenous effector controlling RTK signaling.

### N-WASP recruitment to membrane topology affects Grb2 membrane dwelling

Membrane topology can be modified by curvature inducing proteins (CIPs), which contribute to curvature generation and can also act as membrane curvature sensors [97]. The CIP IRSp53 contains an I-BAR domain, which can interact with N-WASP, which can in turn interact with Grb2 through its Src homology-3 (SH3) domains (Fig. 6E) [98, 99]. N-WASP is an activator of Arp-2/3 complex which drives actin polymerization and exists predominately in a protein complex with WIP (WASP-interacting proteins) [98]. Additionally, membrane curvature regulates the binding affinity of CIPs, which can promote Grb2 recruitment to the membrane and its spatial stabilization (Fig. 6E, left). We thus hypothesized that curved protrusion sites favor the recruitment of CIPs which in turn can recruit N-WASP to the plasma membrane close to RTKs, subsequently these membrane-localized N-WASP interact with Grb2 recruited at the RTKs which could contribute to longer Grb2 dwell times (Fig. 6E, left).

To characterize the dependence of CIP recruitment on membrane curvature, we used dynamically triangulated Monte Carlo simulations [100–102]. Using this framework, we generated membrane topologies defined by three spatial regions, protrusion, annulus, and basal (Fig. 6E top), and computed the excess chemical potential (or the free energy to add/recruit a protein) in each of these regions. Lower value of the excess chemical potential signifies a more favorable recruitment of CIP to a given membrane location. Our free energy calculations showed that the protrusion and annulus regions are preferable sites for CIP localization than surrounding low-curvature basal regions (Fig. 6E, right). Thus, our model predicted that CIPs (such as those containing I-BAR domain) should be favorably recruited to regions with high curvature membrane protrusion sites. This in turn implies that N-WASP would be preferentially localized to CIP-enriched membrane protrusion sites.

To directly analyze adhesion-dependent membrane recruitment of N-WASP, we performed immunostaining for N-WASP followed by TIRF imaging on soft and stiff substrates. We observed an 80% higher recruitment of N-WASP in the cells plated on the stiff substrates compared to the soft substrates (Fig. 6F), without any effect on the total N-WASP levels in the cells (P=0.13; Fig. S13). We also observed the formation of N-WASP puncta in the cells plated on the stiff substrates (inset of stiff in Fig. 6F), which could be regions where Grb2 gets recruited and has an extended dwell time (Fig. 6F). This is in line with our theoretical model, which suggests recruitment of N-WASP to CIP-enriched protrusive areas on the membrane. The diminishment of N-WASP at the membrane is, therefore, predicted to coincide with shortening Grb2 dwell times.

Membrane curvature has been shown to depend on membrane lipid composition, specifically the amount of cholesterol, and disrupting this composition can affect membrane curvature [103, 104]. We, therefore, treated MECs with methyl-β-cyclodextrin (MβCD), an oligosaccharide which depletes cholesterol from the membrane, thus affecting membrane topology [105, 106]. SAIM analysis showed that upon cholesterol depletion with MβCD-dependent cholesterol depletion led to a loss in membrane topology, with a 40% decrease in the FWHM of membrane height (P<0.0001; Fig. 6C, D), consistent with depletion of membrane curvature and a loss of protrusive sites (Fig. 6E, F). However, no reduction in adhesion area was observed with MβCD treatment (Fig. S12.). Thus, MβCD treatment serves as an adhesion-independent way of modulating membrane topology, which can be used to study its effect on N-WASP recruitment and Grb2 membrane dwelling. TIRF imaging of N-WASP immunostained MECs on stiff substrates on MβCD treatment showed a 29% decrease in N-WASP membrane recruitment (P<0.001; Fig. 6D), concurrent with the loss of membrane curvature and loss of CIP from the membrane. Next, we measured the molecular binding dwell time of Grb2 upon cholesterol depletion. Upon cholesterol depletion, the higher Grb2 dwell times that extended up to tens of seconds on stiff substrates were lost (Fig 6G-H). These data further suggest that clustering of Grb2 depends on membrane curvature-dependent N-WASP recruitment, and MβCD-mediated loss of membrane curvature leads to a loss of long-dwelling Grb2 clusters. Thus, our data suggests that the crosstalk between integrin and RTK signaling is likely mediated by membrane topology, which acts as an indirect effector between focal adhesions and RTKs. Thus, ECM stiffness-mediated membrane curvature constrains EGFR into regions with membrane protrusions, leading to Grb2 recruitment and immobilization. These recruited, immobile Grb2 proteins thereafter have the potential to form signaling hubs, where we observed longer dwell time of Grb2, that would likely prolong Ras signaling to increase ERK and PI3Kinase activity.

## Discussion

Many cancers initially respond to tyrosine kinase inhibitor (TKI) therapies, but resistance frequently develops and is attributed to a subset of cells acquiring novel RTK mutations [34, 35, 107]. These dormant cells evade therapeutic intervention and immunological surveillance by lowering expression of proliferation markers that are frequently targeted [34, 35, 108]. Thus, signal amplification in response to microenvironmental changes during cancer progression could promote disease progression driven by these rare cells. Here, we evaluated the known crosstalk between integrins and RTKs [15–17] but raise the additional possibility that resistance does not require acquisition of a mutation or activation of an alternative pathway. Instead, our findings illustrate how relatively modest changes in the lifetime or spatial organization of signaling assemblies can potentiate downstream signaling independent of receptor mutations. We measured the immediate upstream activation of the epidermal growth factor receptor (EGFR) based on docking of Grb2 molecules to phospho-tyrosines. Recruitment of Grb2 occurred across the adherent membrane, suggesting that signal propagation could occur without close proximity to adhesion sites or an increase in receptor expression. Long-dwelling Grb2 events were a small fraction of all events and argue that a handful of receptors can drive signal propagation. Our results suggest that this may be explained by the potency of single SOS molecules that are stably activated at the membrane and achieve a highly processive catalytic state [27, 28, 79]. This phenotype creates a mechanism by which subtle changes in RTK activation, primed by substrate stiffness, are coupled to tuning RTK signaling at all scales.

Integrins mechanically couple the extracellular matrix to the cortical cytoskeleton and organize focal adhesions containing FAK, Src and other signaling components, whereas RTKs such as EGFR provide chemically specific information about extracellular growth factors. These systems are nevertheless strongly interconnected: EGFR activity contributes to rigidity sensing, including Src-dependent EGFR activation during integrin-mediated mechanical probing of the matrix [3], and membrane-associated assemblies can provide physical sites at which integrin and RTK pathways converge. Importantly, receptor crosstalk can also emerge through competition for limiting downstream molecules. Single-molecule measurements have demonstrated that activated EGFR and EphA2 compete for Grb2 and SOS, establishing a hierarchy between receptor inputs based in part on the availability of shared signaling components [4]. Our findings suggest that the robust signaling from a few potent signaling hubs could overcome the competition for signaling adaptors. Accordingly, our data indicate that the physical environment can alter the kinetic availability of signaling effectors while providing strong responses when the effectors are successfully engaged.

Our results support a model in which the plasma membrane acts as an information-processing surface that integrates growth factor signaling tuned by the physical properties of the extracellular microenvironment. The kinetics of individual Grb2 interactions provide an additional dimension through which cells encode information about their surroundings. ECM stiffness-mediated membrane curvature could constrain EGFR into regions with membrane protrusions to enhance Grb2 engagement. These sites would form signaling hubs to facilitate prolonged Ras signaling. The recent development of covalent inhibitors that directly target KRAS4B variants provides a compelling example of why signaling needs to be considered within the physical context of the plasma membrane. Despite the excitement in generating mutant-specific inhibition, acquired resistance still occurs and is not always ascribed to new mutations [109, 110]. Thus, covalently eliminating the activity of one oncogenic Ras species does not necessarily eliminate the capacity of the membrane signaling network to generate Ras activity. The effectiveness and durability of covalent Ras inhibitors may, consequently, depend on events spanning molecular-to-tissue scales based on the kinetics of individual adaptor complexes, through integrin–RTK crosstalk at the membrane, and the stiffness and architecture of the tumor microenvironment. This complexity argues for therapeutic strategies that constrain the adaptive signaling landscape in addition to directly targeting the oncogenic Ras molecule.

Future work is warranted to determine the relative contribution of different proteins and membrane topologies in N-WASP-mediated Grb2 dwell times. Here we used cholesterol depletion as a way to disrupt membrane curvature and observed the depletion of Grb2 long dwells. The computational framework of dynamically triangulated membrane Monte Carlo coupled with free energy methods could also be used to investigate how change in bending rigidity effects curvature-inducing protein recruitment [111]. Therefore, we propose that the stiffness-dependent long-dwelling of Grb2 to the membrane is membrane protrusion mediated, which could be acting downstream of actin polymerization, controlled by integrin signaling [81, 112]. Since membrane composition contributes to topography and availability of lipid signaling molecules like PIP_2_, a mechanistic understanding of the interplay of these components may require new tools. This could be especially important in understanding the roles for lipids, biomolecular condensates, or other machineries in effecting the long-range communication between adhesion and RTK signaling modules.

More broadly, our observations emphasize the importance of understanding signaling as a continuum of information processing across length and time scales. At the smallest scale, individual Grb2 molecules bind membrane-associated receptors for fractions of a second to seconds, while rare stabilized signaling complexes can persist for tens of seconds and disproportionately drive MAPK signaling [1,2]. These molecular events occur within membrane domains and their collective output regulates ERK activity, cytoskeletal organization and cell behavior across tens of micrometers. We therefore propose that environmental sensing is not a pathway operating alongside growth-factor signaling, but a mechanism that continuously reshapes the membrane landscape through which receptor information must pass. In this framework, the biological importance of an input cannot be inferred solely from its abundance: a small number of appropriately positioned and sufficiently long-lived molecular events can be amplified across successive spatial and temporal scales to reshape a global cellular signaling program. Resolving these connections from single molecules to cells and ultimately tissues will be essential for understanding how apparently weak signaling inputs produce robust phenotypes and how the tumor microenvironment enables cancer cells to adapt to targeted therapy.

## Materials and Methods

### Animals and Animal care

All animal husbandry and experimental procedures were performed at the UCSF Parnassus Laboratory Animal Resource Center (LARC) facilities under the Institutional Animal Care Use Committee (IACUC)-approved protocol #AN207099. All experiments used female mice. FVB/NJ (Strain# 001800), PyMT (FVB/N-Tg (MMTV-PyVT)634Mul/J, Strain# 002374) and MMTV-Cre (FVB/N-Tg(MMTV-cre)4Mam/J, Strain# 003553) mice were purchased from The Jackson Laboratory and described before [66, 113]. Genetic backgrounds were confirmed by genetic monitoring (TransnetYX). To inhibit lysyl oxidase (LOX) activity, PyMT mice received drinking water supplemented with 0.5 mg/mL β-aminopropionitrile (BAPN, TCI, Cat# A0796) beginning at four weeks of age. Mice were sacrificed at thirteen weeks of age for tissue analysis.

### Immunofluorescent staining

For phospho-FAK and phospho-ERK staining of tissues, OCT-embedded frozen mammary tumor tissue sections were used. Sections were fixed in 4% paraformaldehyde (PFA), and bleached to reduce autofluorescence using 4.5% (wt/vol) hydrogen peroxide and 20 mM NaOH in PBS and placed under LED lights twice for 30 mins [114]. Tissues were then permeabilized with 0.1% Triton X-100 in PBS and blocked with 5 % BSA and 5 % goat serum in PBS. Samples were incubated with primary antibodies (phospho-Tyr397 FAK: Invitrogen Cat #44-625G; or phospho-p44/42 MAPK (ERK1/2)(T202/Y204): Cell Signaling Technology, Cat #9101; along with Cytokeratin 8+18 (pan-CK): Fitzgerald, Cat # 20R-CP004; Cytokeratin 5: Fitzgerald, Cat # 20R-CP003) overnight at 4°C, washed and then incubated for 1 hr at RT with species-specific secondary antibodies (Thermo Fisher Scientific, Cat #11001, #A11011, #A11073) conjugated to fluorophores and CNA35-GFP (Addgene, Plasmid #61603) where specified. Cell nuclei were counterstained with 4,6-diamidino-2-phenylindole (DAPI, Thermo Fisher Scientific, Cat #D1306). Images were acquired with Nikon SoRa spinning disk microscope equipped with a 40x water-immersion objective. Image analysis was performed in ImageJ. For phospho-ERK quantification, percentage positive cells from pan-CK positive cells were counted. For phospho-FAK quantification, pan-CK positive cells were analyzed for intensity of pFAK stain and background subtracted.

For immunostaining for cells seeded on glass or silicone gels, adhered cells were fixed with 4% paraformaldehyde for 15 min at 37℃, permeabilized in 0.1% Triton X-100 for 20 min, blocked with 1% BSA for 1 hr, and incubated with primary antibody overnight at 4℃. Subsequently, cells were washed with PBS and incubated with a fluorescently labelled secondary antibody and phalloidin-Alexa 561 (Thermo Fisher Scientific, Cat #; 1:200) for labelling actin for 2 hr at room temperature. The coverslips were then washed with PBS and stored at 4℃ till imaging.

### Immunohistochemical staining

For phospho-AKT substrate staining, mammary tumors were collected, formalin-fixed and paraffin embedded (FFPE) and processed for histology by cross-sectional profile cuts. FFPE tissue sections were dewaxed and rehydrated through xylene and graded alcohols. Antigen retrieval was performed by boiling sections in 10 mM citrate buffer. The tissue sections were then blocked with 5% BSA and 5% goat serum in PBS and incubated with primary antibodies (phospho-AKT substrate (RXXS*/T*) (110B7E): Cell Signaling Technology, Cat #9614) overnight at 4℃. Tissue sections were then washed with PBS and incubated for 1 h with horseradish peroxidase (HRP)-conjugated secondary antibodies (ImmPRESS HRP Goat Anti-Rabbit IgG Polymer Detection Kit, Vector Laboratories, Cat# MP-7451). Stain development was performed using the Tyramide Signal Amplification method, using CF488A Tyramide substrate (Biotium, Cat #92171), and nuclei were counterstained with DAPI. For phospho-AKT substrate quantification, stained tissues were analyzed for mean intensity of the stain, and background subtracted.

### STIFMap image analysis

STIFMap images were generated as previously described [46]. Briefly, immunofluorescent-stained images of CNA35-GFP and DAPI were used to generate STIFMaps using five independently trained models. The predicted elastic modulus was pseudocolored using the ‘Fire’ colormap, and the top 10% of stiffness values for each tissue were used to calculate the mean predicted elastic modulus in pascals.

### Cell Culture and Transfection

MCF10A cells were maintained in growth medium (DMEM/F12 Ham’s mixture (Invitrogen), 5% horse serum (Invitrogen), 20 ng/mL EGF (PeproTech), 0.5 ug/mL hydrocortisone (Sigma) (Sigma), 100 ng/mL cholera toxin (Sigma, Cat 3C8052), 10 ug/mL insulin (Sigma) and 1x Pen/Strep) at 37°C in a humidified atmosphere with 5% carbon dioxide. HEK293T cells were cultured in DMEM (Genesee Scientific) supplemented with 10% FBS (Gemini Bio-Products). Cells were regularly tested for mycoplasma contamination.

Cells were transfected with 1 ug DNA per 1 million cells in serum-free media using Lipofectamine 3000 (Invitrogen). Plasmid solutions were added dropwise to cells and incubated for 4 hours before replacement with fresh media. Cells were allowed to express the transfected constructs for 24-36 hours prior to imaging. For knockdown or inducible expression, cells were incubated with the appropriate reagents diluted in growth medium. To generate stable cell lines, cells were retrovirally transduced with HEK293T cell-derived supernatants. HEK293T cells were transfected with appropriate vectors using PEI/Lipofectamine 2000 and allowed to express viral particles for 48-72 hours.

Cells were seeded at a density of 20,000 cells/mL on fibronectin-conjugated substrates (glass: ethanol-flamed 25 mm coverslips coated with 10 mg/mL fibronectin; and silicone gels) to have single cells that were well separated. Cells were plated for a minimum of 15 hours prior to live imaging or fixation. In preparation for live imaging, cells were incubated in phenol-free medium to reduce background signal. Cells were starved for 4 hours in phenol-free, serum-free DMEM/F12 prior to imaging. Cells were stimulated with 100 ng/mL EGF, either unlabeled or labeled with AlexaFluor 647 (Thermo Fisher), in phenol-free, serum-free DMEM/F12 media + 0.1% (wt/vol) BSA. Live cell imaging was achieved in an environmental chamber at 37°C and 5% CO_2_.

### 3D spheroid culture

Collagen type 1/reconstituted basement membrane (Col/rBM) hydrogels were prepared as previously described. Briefly, 2 mg/mL rat tail Col1 (Corning, Cat# 354249) was incubated with 0.1% acetic acid (non-crosslinked; soft) or 0.1% acetic acid with 500 mM L-ribose (Chem Impex, Cat# 28127) (crosslinked; stiff) for 10 days. The Col1 mixtures were then combined with BM extract (R&D Systems, Cat# 3532-005-02) at a final concentration of 20%. MCF10A spheroids were resuspended in the soft and stiff collagen/BM mixtures, plated as a 100 μL drop, and allowed to solidify for 30 minutes at room temperature, then for 30 minutes at 37°C, followed by the addition of complete medium.

### Western Blot

Following starvation and stimulation with EGF, 3D spheroids were lysed in SDS lysis buffer (2% SDS, 50mM Tris-HCl (pH 8), and 10mM EDTA) supplemented with protease and phosphatase inhibitors, and lysates were collected at multiple time points and used in immunoblotting. The lysate was sonicated by 10 short bursts with a stainless steel probe sonicator before boiling at 95°C for 5 min. Samples were centrifuged at 13,000 g for 10 min, resolved on SDS-PAGE gels, and transferred onto polyvinylidene difluoride (PVDF) membranes (BioRad). The membranes were blocked with 5% BSA for phosphorylated antibodies or 5% milk otherwise in TBST and probed with the indicated primary antibodies in 5% BSA/TBST, and a 1:10,000 dilution of anti-rabbit or anti-mouse horseradish peroxidase-conjugated secondary antibodies in 5% BSA/TBST. Blots were developed using WesternBright Quantum (Advansta) and visualized on PXI 6 Touch (Syngene). The band intensity of all Western blots was quantified using Fiji software. Antibodies used for western blotting: phospho-p44/42 MAPK (ERK1/2) (T202/Y204): Cell Signaling Technology, Cat. #9101; p44/42 MAPK (ERK1/2)= Cell Signaling Technology, Cat. #9102; GAPDH: Cell Signaling Technology, Cat. #2118; Phospho-EGFR (Tyr1068) (D7A5): Cell Signaling Technology, Cat. #3777; EGFR: Cell Signaling Technology, Cat. #4267; HRP-conjugated goat anti-rabbit: Advansta, Cat. #R-05072-500; HRP-conjugated goat anti-mouse: Advansta, Cat. #R-05071-500.

### Drug treatments

Focal adhesion kinase activity was impaired using the selective, small molecule, FAK Inhibitor 14 (Tocris, Cat no. 3414) at a 60 nM concentration. methyl-β-cyclodextrin (MβCD) treatment (Thermo Scientific, Cat. #J66847.06). Knockdown of wild type beta 1 integrin was accomplished by a 5-day treatment with 100 mM Isopropyl β-d-1-thiogalactopyranoside (IPTG) (Sigma, Cat. #10724815001) to induce the shRNA construct. The V737N integrin mutant was expressed by 24 hours of inducible expression with 100 ng/mL of doxycycline (Alfa Aesar, Cat. #J60579).

### Silicone gel preparation

Silicone gels were deposited onto NaOH-cleaned glass-bottom chambers (Cellvis) or on 25 mm glass coverslips by spin coating, as previously described [89]. Briefly, polymers (Corning 52-276 Parts A and B) were mixed at fixed ratios (1.6:1 corresponding to soft and 1:1.6 corresponding to stiff) and degassed. The gel solution was spin-coated (VTC-100, MTI Corporation) onto the glass substrate prior to 3 hours of curing in an 80°C incubator. Under aseptic conditions, gel surfaces were activated with 0.5% (3-aminopropyl)trimethyloxysilane (ATPES; ACROS Organics, Cat. #313251000) in 95% ethanol for 5 minutes. Following activation, 100 µg/mL 1-ethyl-3-(3-dimethylamino-propyl)carbodiimide (EDC; Chem-Impex Int’l Inc., Cat. #00050) and 10 mg/mL fibronectin (Millipore; Cat. #FC010) were incubated with the gel for 30-60 minutes. Once gels were activated, derivatized with fibronectin, and washed, cells were immediately plated onto the gel surfaces. Fibronectin labeled with AlexaFluor 569 was used to confirm functionalization by FRAP. In all other experiments, non-fluorescent fibronectin was used.

### Atomic force microscopy (AFM) of tissues

AFM measurements were conducted using an MFP-3D BIO inverted optical AFM (Asylum Research) mounted on a Nikon TE2000-U inverted fluorescent microscope (Melville), stabilized on a vibration-isolation table (Herzan, TS-150). Indentations were performed using silicon nitride cantilevers with a nominal spring constant of 0.06 N/m, fitted with a 5 μm-diameter spherical borosilicate glass tip (Novascan Technologies, PT.GS). Cantilevers were calibrated using the thermal fluctuation method. OCT-embedded frozen tissue sections of 20 μm thickness were thawed and equilibrated to room temperature before immersion in PBS containing protease and phosphatase (GenDEPOT, Cat# P3100 and #P3200) inhibitors. Tissue specimens were indented at a 2 μm/s loading rate. The Young’s elastic modulus was calculated by fitting force curves using the Hertz model, assuming a Poisson ratio (υ) of 0.5.

### Shear rheology of Col1/rBM and silicone gels

Rheological measurements of Col1/BM and silicone gels were performed using a stress-controlled AR2000EX rheometer (TA Instruments), as previously described. Col1/rBM hydrogels or silicone gels were excised using an 8 mm biopsy punch (Kai Medical, Cat# BP-80F) and then placed directly onto the lower Peltier plate and brought into contact with a parallel plate geometry, forming a disk-shaped sample. An oscillatory strain sweep from 0.1% to 1% was performed at a constant angular frequency of 1 rad/s. Storage (G′) modulus reached equilibrium during the sweep, and values reported reflect measurements at 1% strain. The elastic modulus (E) was calculated from the storage modulus using the equation: E = 2G′(1 + υ), assuming a Poisson’s ratio (υ) of 0.5 for both the hydrogels and silicone gels.

### DNA Constructs

The gene sequence encoding photo-convertible mEos3.2 was cloned into a pCDNA3.1 mammalian expression vector containing a CMV promoter. mEos3.2 was amplified from the mEos3.2-C1 plasmid submitted to Addgene (#54550) by Michael Davidson (Florida State University) using standard PCR methods. mEos3.2 is a derivative of mEos2 that contains the following point mutations: I102N, H158E, Y189A (Excitation=507/572 nm; Emission= 516/580nm). Using restriction-free cloning (van den Ent & Lowe, 2006), DNA sequences encoding human Grb2 (1-217aa) or the C-terminal hypervariable sequence derived from human K-Ras isoform 2B (-GSKMSKDGKKKKKKSKTKCVIM*) were inserted at the 3’ end of the open reading frame of mEos3.2. The resulting expression constructs include plasma membrane-targeted mEos3.2-CAAX and mEos3.2-GGG-Grb2(1-217aa), with a (Glycine)x3 linker between the mEos3.2 and Grb2 domains. Inducible V737N β1 integrin construct has been previously described [115] as puromycin lentiviral transfer vector expressing 3x myc tagged V737N β1 integrin using the tetracycline rtTA2(S)-M2 (Urlinger et al., 2000) inducible promoter for 24 h with doxycycline.

### Microscopy

Imaging was performed on an inverted microscope (Nikon Eclipse Ti-E; Technical Instruments, Burlingame, CA) equipped with a motorized stage (Applied Scientific Instrumentation MS-2000, Eugene, OR). Illumination was provided by either a mercury lamp (IntensiLight; Nikon C-HGFIE) or a custom-built laser launch (Solamere Technology Group, Inc., Salt Lake City, UT) equipped with 405, 488, 560, and 640 nm diode lasers (Coherent OBIS, Santa Clara, CA). RICM excitation was filtered using a 50/50 beamsplitter. For TIRF imaging, laser excitation was reflected using a dichroic mirror (SPECS of Quad Cube) to a 100x objective lens (Nikon, 1.47 NA; Technical Instruments, Burlingame, CA). Emission was collected using the appropriate filters (ET525/50M, ET600/50M, ET700/75M; Chroma Technology Corp., Bellows Falls, VT) on an EM-CCD device (iXon 897DU; Andor Inc., South Windsor, CT). All imaging hardware was controlled using the MicroManager software package [116].

### Imaging Conditions

Stochastic switching of mEos3.2 constructs was achieved using a pulse of 405 nm light TIRF angles were optimized for each sample to selectively illuminate the basement membrane. Typical acquisitions used 50 millisecond exposure times and a 0 second interval (streaming). Acquisitions were set as 4000-frame cycles with 1000-frames of single molecule mEos3.2 imaging as 561 nm separated by single images of RICM and fluorescent EGF by 640 nm excitation. These settings resulted in datasets lasting approximately 4 minutes and 10 seconds. For imaging stimulated cells, image acquisitions were initiated immediately after addition of fluorescently-labeled EGF.

### Image processing

Image stacks were parsed into 1000-frame segments. Single molecule fitting and tracking were performed using the plugin Octane for ImageJ, with an adaptive threshold, max displacement of 2 pixels, and no frame skips. Trajectories were exported to MATLAB (R2023b) for additional processing. Dwell times were calculated as the number of frames each particle was tracked for. A lower threshold cutoff of 5 frames (250 ms) was used. To plot dwell times, the plotTracks function from msdanalyzer package (https://tinevez.github.io/msdanalyzer/) was used, along with a defined cell boundary. Each track was normalized into a *jet* colormap according to their track duration and categorized into pre-defined bins (e.g., 0-2 s, 2-5 s, >5 s for Figure 2F). Dwell time probability distributions were plotted on a log scale as a function of dwell time in seconds.

For spatial analysis of SPT tracks, tracks with a duration greater than 2 seconds were condensed into the center of mass (COM) of the entire track, COMs outside the cell boundary were eliminated, and distances of COMs to cell edge were calculated. Concentric bin boundaries of the same area were generated and COMs were classified into the bins based on their distance from the edge. Mean track duration and probability of long events (tracks in the bin divided by total number of tracks) were calculated and plotted as a function of bins (center to periphery). For spatial analysis of adhesions, pFAK images were thresholded into adhesion masks and cell boundaries were created using phalloidin. Cells were divided into 10 area-equal bins using the *quantile* function in MATLAB, and each track COM was assigned a bin depending on its position. Large adhesions were defined as size > 0.5 µm^2^, and the probability of large adhesions was calculated by number of large adhesions in the bin divided by total number of large adhesions in the cell.

To quantify cell spreading and circularity, fluorescence images were analyzed with ImageJ software. F-actin staining by phalloidin was used to mark the cell boundary, to determine the projected area and circularity of the cells. Phospho-FAK staining was used to calculate the adhesion area and adhesion length. A background size threshold of 0.01 µm^2^ was set for identifying adhesions. To quantify N-WASP fluorescent intensity at the membrane, TIRF images of actin staining were used to create a cell boundary, N-WASP intensity inside the boundary was averaged, and then background subtracted. All values were normalized to the average of the soft condition.

The number of Grb2 molecules recruited in response to stimulation was estimated based on new localizations per unit time. Data were measured after converting a substantial fraction of Grb2-mEos3.2 molecules to minimize undercounting. Correlation analysis of SOS-GFP and Paxillin-mCherry was performed using the coloc 2 plugin on ImageJ, within the cell boundary. The Pearson’s coefficient was calculated and depicted.

### Scanning Angle Interference Microscopy

Silicon oxide doped silicon wafers (Addison Engineering) substrates were prepared by ATPES (ACROS Organics, Cat. #313251000) and glutaraldehyde activation, followed by conjugation with 10 μg/ml fibronectin (Millipore). Cells ectopically expressing membrane bound-CAAX tagged with mEmerald fluorophore were plated overnight on the fibronectin-coated wafers, fixed and imaged [96, 117]. The wafers were inverted on a drop of PBS, weighed down using a lock nut, and imaged using a 488 nm laser with an incident angle of excitation light from -48° to +48° with a one-degree sampling rate. Z-positions were localized with custom algorithms [96] and membrane height values were extracted and plotted as a 3D image using the *mesh* function with a *turbo* colormap in MATLAB (R2023b). Membrane height values inside the cell boundary were extracted, and a histogram of these values were used to calculate the full width half maxima (FWHM) for each cell.

### Flow cytometry

For the quantification of EGFR surface density, cells were plated on polyacrylamide gels of soft (400 Pa) or stiff (8.4 kPa) elastic moduli. Cells were detached from the gels using Accutase and incubated with the fluorochrome-conjugated secondary antibodies (Invitrogen, Cat #A11008) against EGFR (Cell Signaling, Cat #4267S) for 30 minutes on ice. Cells were pelleted, washed and suspended in PBS, strained through 40 µm filters, and analyzed using a Becton Dickinson LSR II or Fortessa with the channels corresponding to the fluorochromes. All flow cytometry data was collected using FACS Diva and analyzed and plotted using FloJo and Graphpad Prism v6.0.

### Biophysical Modeling

The description of modeling methods used for the free energy landscape for protein recruitment on the cell membranes are detailed in this section.

#### Mesoscale membrane model

The membrane simulations consist of a membrane patch, modeled as a thin continuum elastic surface, evolved with Monte Carlo according to the Canham-Helfrich Hamiltonian. This method is adapted from techniques described in [100]. Briefly, the membrane patch is discretized into a set of interconnected triangles with each triangle representing a patch of a bilayer whose size is much smaller than the persistence length of the bilayer. The discretized surface consists of *N* vertices, each of characteristic size *a*_0_, interlinked by *L* links that form *T* triangles. The thermodynamic behavior of the membrane is well captured by the elastic energy functional known as the Canham-Helfrich Hamiltonian [118, 119].

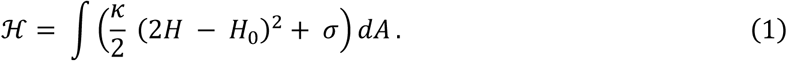

Here, *κ* and *σ* are the bending rigidity and the surface tension of the membrane. For a triangulated membrane patch, the discretized form of the Hamiltonian in Eq. (1) is

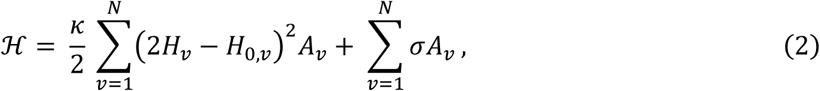

Where *H_v_* and *A_v_* are the mean curvature and the area associated with vertex *v*, computed using the methodology introduced by Ramakrishnan et al. [101]. *H*_0,*v*_ is the spontaneous curvature associated with vertex *v*, imposed on the membrane by a curvature modulating factor that can be modified to model different biophysical processes.

For generation of membrane protrusion, a spontaneous curvature field is assumed to be a circular collection of vertices of a prescribed spontaneous curvature with a nearest-neighbor (Ising model-like) potential keeping these vertices together. This circular region of spontaneous curvature is meant to approximate the effects of a large complex protein-assembly involved in pushing out a protrusion [120]. The spontaneous curvature field for generating the membrane protrusion is applied at the center of the membrane,

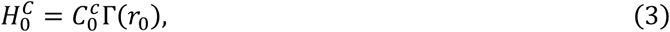

where Γ(*r*_0_) is a function that is unity within a circular domain of radius *r*_0_ and zero otherwise, and 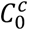 is the magnitude of imposed curvature.

For a single curvature-induing protein, a spontaneous curvature field is assumed to be the form of a Gaussian function with a pre-factor 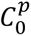 (peak spontaneous curvature), and a variance proportional to *ε*^2^ (field spatial extent),

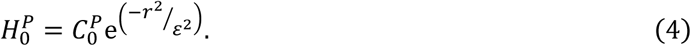

This approach of treating the effect of the curvature-inducing proteins containing BAR and ENTH domains as a curvature field in the continuum field formulation has been utilized in prior studies [100]. A protein field can is associated with a unique vertex and a vertex can accommodate at most one protein field. The contribution of a protein field to the spontaneous curvature associated with a vertex in the vicinity is determined by *r*, the distance of the field from the vertex.

In our simulations, the membrane is modelled with fixed projected area (equal to that spanned by the x–y plane) in a cubic box with periodic boundaries in two-dimensions (i.e. along the x–y plane). The system is evolved using a set of three Monte Carlo moves: the vertex displacement, which simulates thermal undulations; the link flip, which simulates membrane fluidity and allows it to more dynamically remodel, and the protein move, which allows protein movement along the membrane. All three MC moves are accepted with Metropolis acceptance criteria.

The results presented in this study are for a membrane patch with *N* = 900, *κ* = 20 *k_B_T*, and 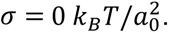 Here, *k_B_* is the Boltzmann Constant, T is the temperature (set to 300 K), and *a*_0_ is the characteristic length scale for the triangulated surface. To generate membrane protrusion, 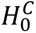 with 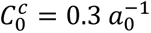 was applied at the center of the membrane that consists of 50 vertices; this corresponds to *r*_0_ = 4.55 *a*_0_. The protrusion region was defined as all 50 vertices with the spontaneous curvature field. The annulus region, a radial extension of the protrusion region, was defined by creating a neighbor list at the point of insertion: if a protrusion region vertex existed within the first 30 closest neighbors of this vertex and the vertex was not a protrusion vertex itself, this was described as the annulus region vertex. Vertices that do not meet either of these criterions were categorized as the basal region. The curvature-inducing proteins inserted in this study had peak spontaneous curvature of 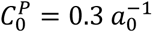 and a variance of 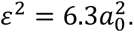.

#### Free energy calculations

Previous studies have coupled the membrane model with computational free energy methods [102, 120]. We employ Widom insertion to compute the excess chemical potential associated with inclusion of a curvature-inducing protein on a membrane. It involves randomly inserting a protein field and recording the difference in the in energy between a system with *n* and *n* + 1 proteins which relates to the excess chemical potential for (*n* + 1)^th^ protein as

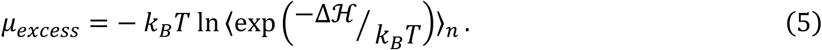

Here, Δℋ is the change in energy calculated from the Canham-Helfrich Hamiltonian upon insertion of that protein and 〈·〉 corresponds to the Boltzmann weighted ensemble average. This technique can also be extended to compute spatially varying chemical potential (inhomogeneous Widom insertion [102, 121], whereby each subsection has its excess chemical potential defined as

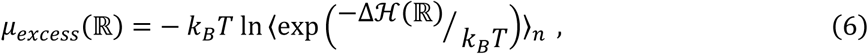

where ℝ defines the spatially distinct region. In the case of triangulated membrane, ℝ corresponds to a collection of vertexes within a specified region. For protrusion simulations, where a circular region induces spontaneous curvature 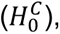, inhomogeneous Widom insertion was done in three spatially distinct regions: protrusion, annulus, and basal regions. This study was conducted with no membrane proteins present (i.e. *n* = 0 and *n* + 1 = 1).

When conducting inhomogeneous Widom calculations, the background spontaneous curvature of the protrusion-field was disregarded in the Widom formula. Curvature-inducing proteins with spontaneous curvature fields 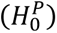 were then randomly inserted and categorized into the appropriate region. In all Widom results shown, simulations were equilibrated for 5 million Monte Carlo steps and sampled with Widom method every 100 steps. Only the last 2.5 million Monte Carlo steps were used when computing the ensemble average to ensure an equilibrated system for sampling. All reported data is averaged over three independent ensembles.

The form of the excess chemical potential of a curvature-inducing protein can be derived from Eq. (1). The energy difference between a state without a curvature-inducing protein and after the addition of the curvature-inducing protein is given by

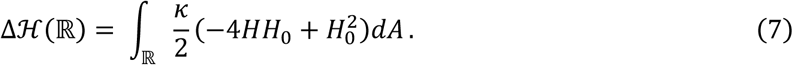

Substituting Eq. (7) in Eq. (5) for excess chemical potential and incorporating the curvature field for a curvature-inducing protein (Eq. (4)), we get

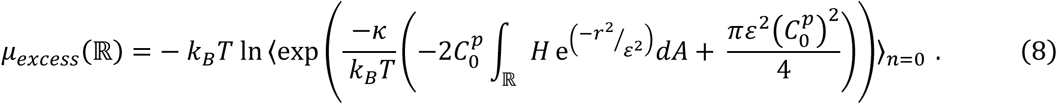

Since the second term in the exponential depends only on constants, this relation can be further simplified to

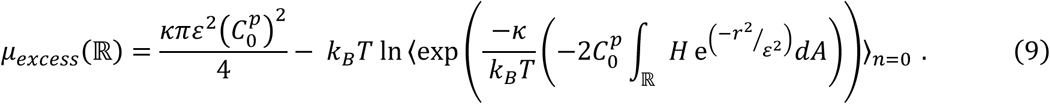

The first term in Eq. (9) is a constant and can be interpreted as the chemical potential to insert a protein on a flat membrane. The second term accounts for contribution of membrane fluctuations at finite temperature to recruitment of a curvature-inducing protein, whereby the localization to membrane sites matching the curvature profile of the curvature-inducing protein increases magnitude of second term, thus reducing *μ_excess_* and providing favorable conditions for curvature-inducing protein recruitment. Supporting this theoretical framework, free energy calculations in dynamically triangulated Monte Carlo simulations (Fig. 6E, right) showed that excess chemical potential for curvature-inducing protein has lower value at the protrusion site as compared to the surrounding annulus and planar basal regions.

### Quantification and statistical analysis

All experiments were repeated at least three times where we measured >5 cells in each repeat. Graphs were plotted and statistical analysis was performed using GraphPad Prism 8. Individual data points were plotted along with the box plots with P-values written on top of the graph. The number of biological replicates (n) is indicated in each figure legend. Data are presented as mean ± SD. A p-value < 0.05 was considered statistically significant.

Data from individual groups was first analyzed for normal distribution using the Shapiro–Wilk normality test. If all groups were normally distributed, a two-tailed t-test was used for 2 groups, or an ordinary one-way analysis of variance (ANOVA), followed by Tukey’s post hoc test, was used for more than 2 groups was performed. If all groups were not normally distributed, the Mann-Whitney test was used for 2 groups, or the Kruskal-Wallis test, followed by Dunn’s post hoc test, was used for more than 2 groups.

## Competing Interests

The authors declare no competing interests.

## Acknowledgements

This work was supported by U.S. National Institutes of Health NCI U01 grant CA202241 to J.G. and V.M.W. and R35CA242447 and Weill Project FEAST to V.M.W., and a Catherine Peachey Breast Cancer Research Grant to STL-N. K.J. is the Lois A. Cinelli Fellow of the Damon Runyon Cancer Research Foundation (DRG-2547-25). We thank FuiBoon Kai (UCSF) and Catherine B. Carbone (UCSF) for help with SAIM, Shelly Kaushik (UCSF) for help with Flow cytometry, Dongmyung Oh (NUS, Singapore) and Guanqing Ou, (UCSF) for helpful discussions.

## Data availability

All data supporting the findings of this study are available from the corresponding authors on reasonable request.

## Contributions

S.T.L.-N., D.T, V.M.W. and J.T.G. conceived the project and designed experiments. S.D.H. and J.N.L. performed cloning and supported *in vitro* experiments. K.J., S.T.L.-N., J.J.Y.L., and D.T. performed *in vitro* experiments and analyses. K.J., M.-K.H. and M.M. performed *in vivo* experiments and analyses. K.P. and R.R. performed mathematical simulations. S.T.L.-N., K.J. and V.M.W. wrote the manuscript. All authors proofread the manuscript.

## Notes

### Competing Interest Statement

The authors have declared no competing interest.

