## Supplemental figures for "Single molecule Grb2 dynamics reveal spatiomechanical modulation of RTK signaling across length scales"

Supplementary Figures

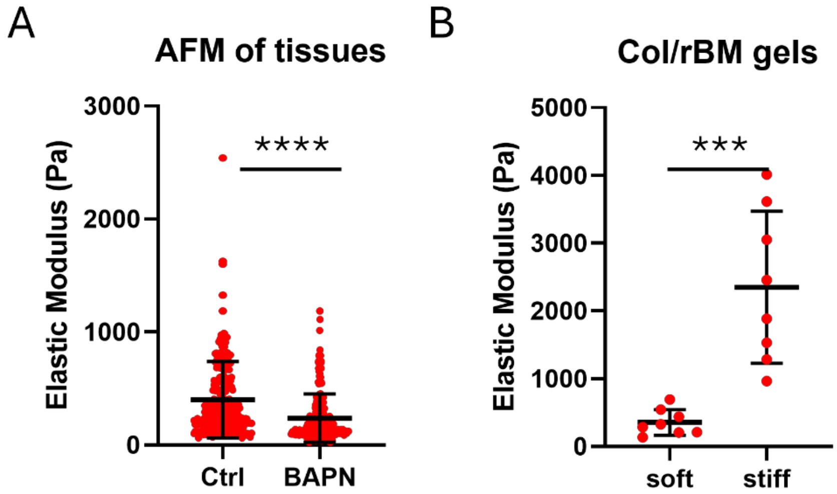


Figure S1. Elastic modulus measurements for tissues and 3D Col/rBM gels. (A) Atomic Force Microscopy (AFM) measurements of MMTV-PyMT vehicle (Ctrl) vs. BAPN-treated mammary tissues. (B) Rheology measurements of Col/rBM with (stiff) or without (soft) ribose crosslinking were used to calculate the elastic modulus of the gels. Data are plotted with biological replicates as scatter points with mean±s.d. Statistical significance was measured by Mann-Whitney test (A) unpaired t-test (B) ***$p < 0.001$, ****$p < 0.0001$.

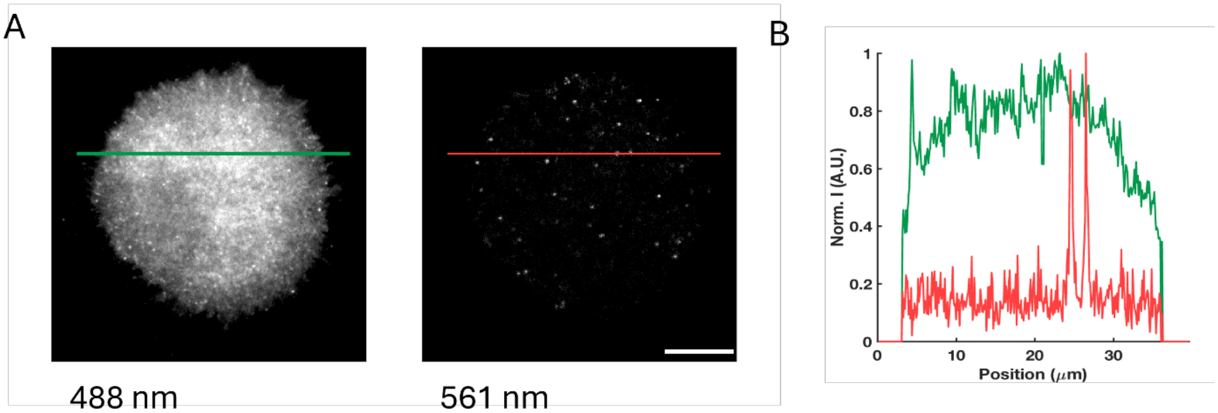


Figure S2. A) MECs expressing mEos3.2-Grb2 imaged in TIRF using 488 excitation (left) and 561 excitation after a short pulse of 405 nm light for photoconversion (right). B) Intensity line scan across TIRF images showing intensities in 488 nm (green) and 561 nm (red) images. Scale bar = 5 μm.

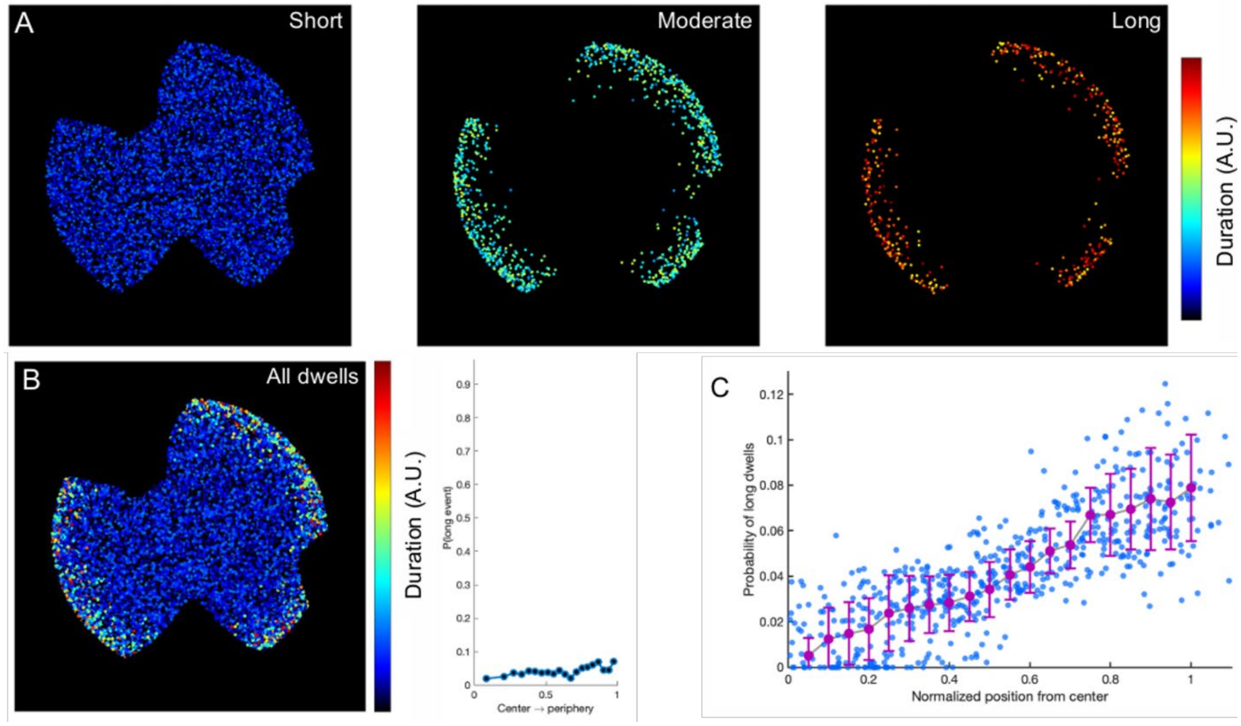


Figure S3. Simulations and spatial analysis for a heterogeneous distribution. A) Dwells were simulated as points with a mean dwell time divided into short, moderate and long tracks. The moderate and long tracks were simulated close to the edge, while short tracks were simulated all across the cell shape. B) A merge of all dwells, along with the spatial analysis of the probability of long dwells as a function of distance from the center to the periphery. C) A plot of probability of long dwells as a function of distance from the cell center to the cell periphery.

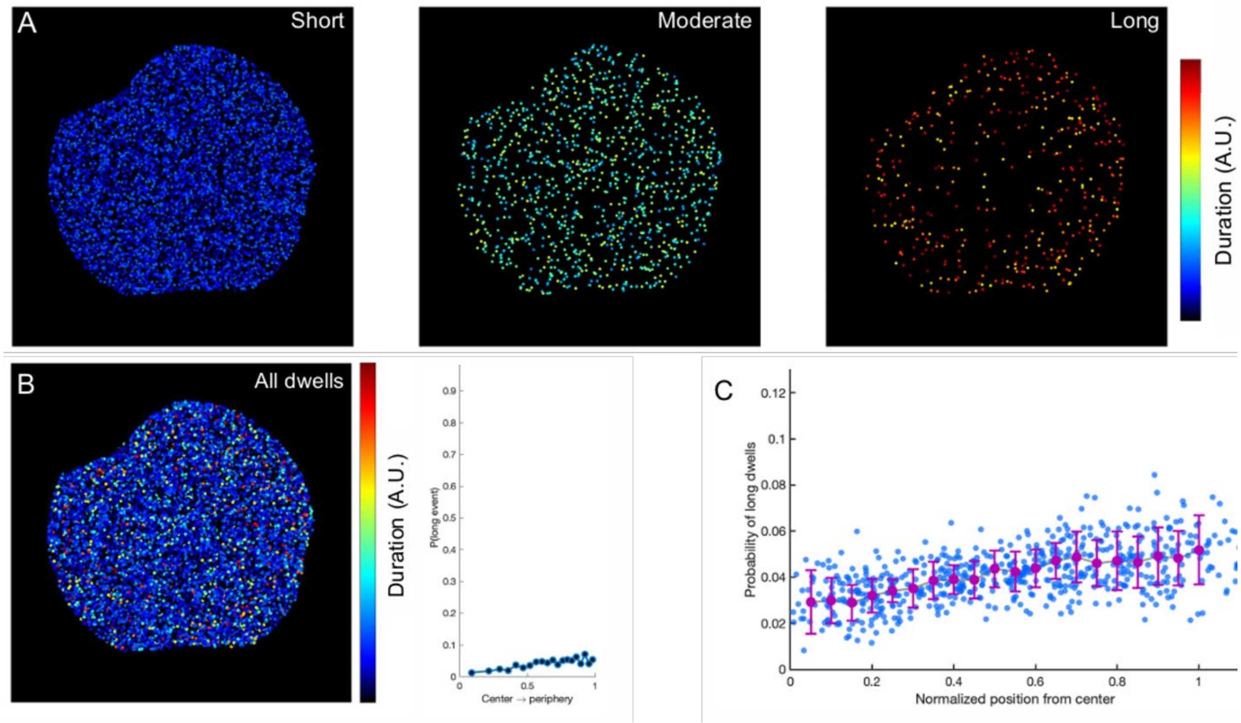


<u>Figure S4.</u> Simulations and spatial analysis for a homogeneous distribution. A) Dwells were simulated as points with a mean dwell time divided into short, moderate and long tracks, all distributed homogenously across the cell membrane. B) A merge of all dwells, along with the spatial analysis of the probability of long dwells as a function of distance from the center to the periphery. C) A plot of probability of long dwells as a function of distance from the cell center to the cell periphery.

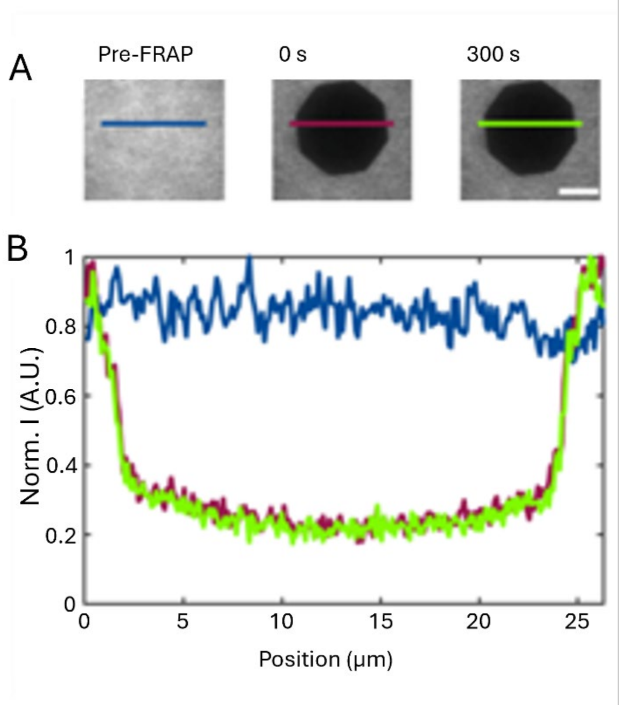


<u>Figure S5.</u> Validation of fibronectin functionalization on silicone gel substrates. (A) Fluorescence Recovery After Photobleaching (FRAP) of fluorescent fibronectin on stiff gel. (B) The intensity line profile of the bleached region. No fluorescence recovery, as shown by the line scan, indicates that fibronectin is present and immobile.

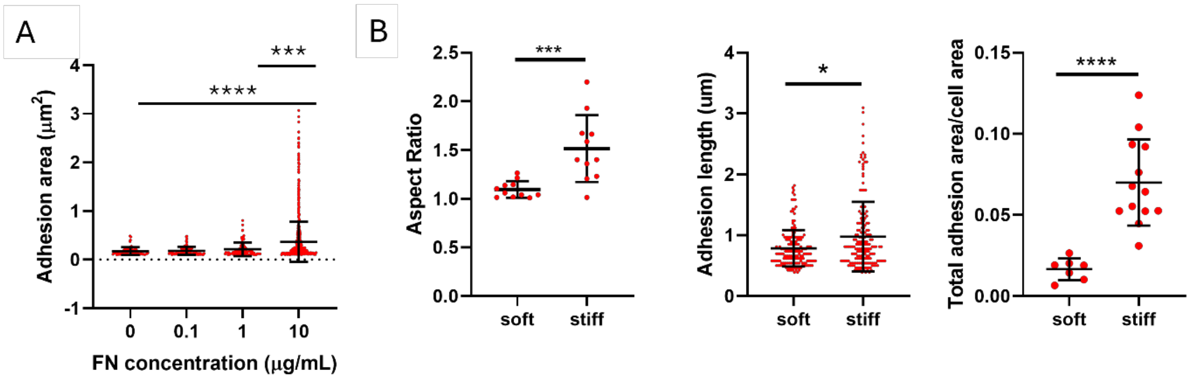


Figure S6. Cell spreading and adhesions on fibronectin and 2D gels. (A) Adhesion area quantification for MEC cells spread on the different fibronectin concentrations (0, 0.1, 1, 10 μg/mL). Data are plotted with cells as scatter points with mean±s.d. Statistical significance was measured by ANOVA followed by the Kruskal-Wallis test; ***$p<0.001$, ****$p < 0.0001$. (B) Quantification of cell aspect ratio, adhesion length, and total adhesion area normalized to cell area for MEC cells spread on fibronectin-coated soft and stiff silicone gels. Data are plotted with cells as scatter points with mean±s.d. Statistical significance was measured by unpaired t-test or Mann-Whitney test; *$p>0.05$, ***$p < 0.001$, ****$p<0.0001$.

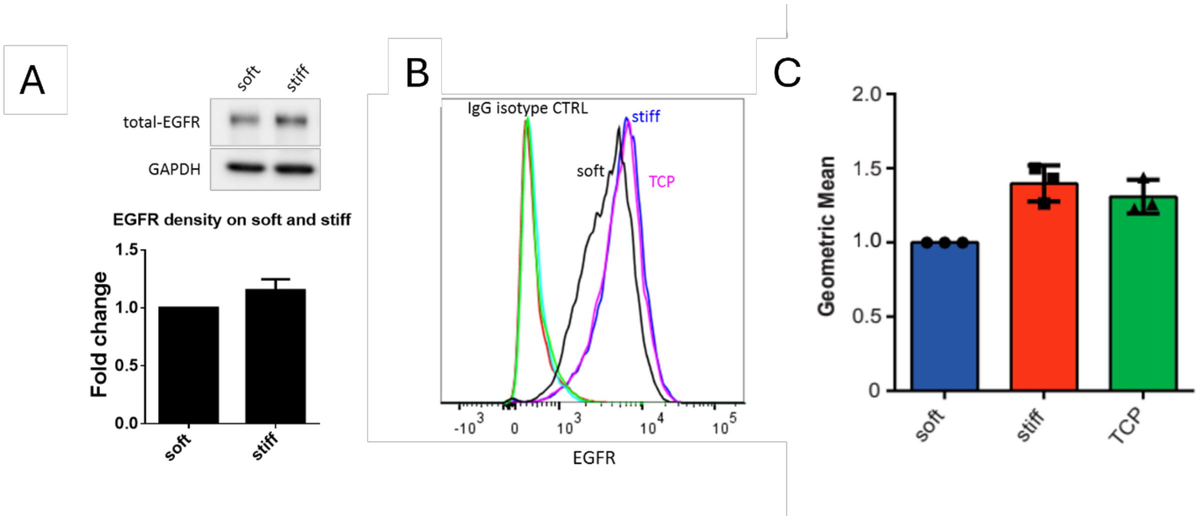


Figure S7. Stiffness does not alter EGFR surface density. (A) Quantification of total EGFR density of MECs on soft and stiff gels by Western Blotting. (B) Quantification of EGFR density of MECs on soft, stiff gels or on TCP by flow cytometry. C) Analysis demonstrate that there is a slight (non-significant) increase in the surface density of EGFR of MECs plated on stiff vs soft gels.

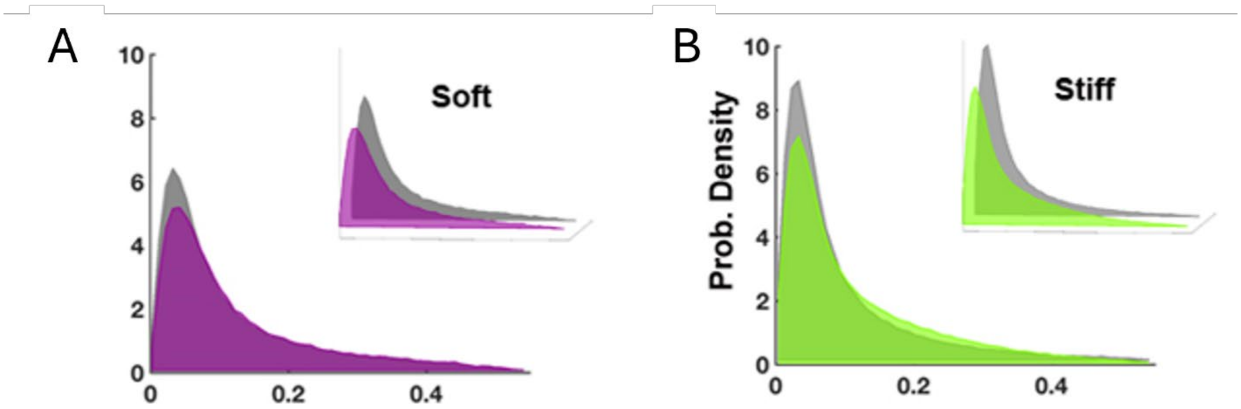


Figure S8. Grb2 mobility is modulated by substrate stiffness. (A, B) Step size distributions for Grb2 in MECs on (A) soft gels, and (B) stiff gels. In each case, the colored distribution represents step sizes under starvation conditions. In each case, the gray distribution indicates step sizes following stimulation with 100 ng/mL EGF. Step size distributions are shown enlarged in 2D with an inset showing the same distributions in 3D. N>5 cells each.

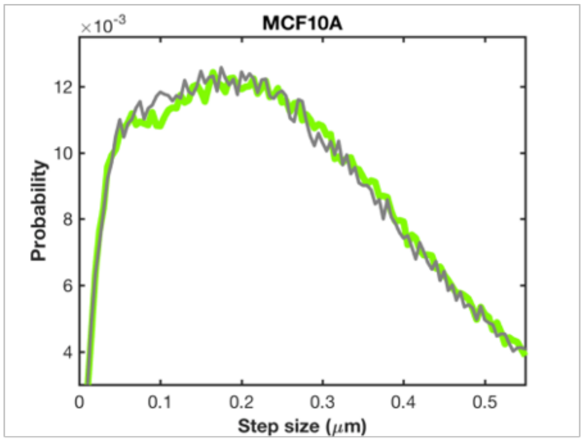


D = 0.02 μm²/s, α = 19%
D = 0.01 μm²/s, α = 21%

Figure S9. Mobility of a membrane anchored construct is unaffected by EGF stimulation. Step size distribution of mEos3.2-CAAX before (green) and after (gray) stimulation with 100 ng/mL EGF shows no change in behavior.

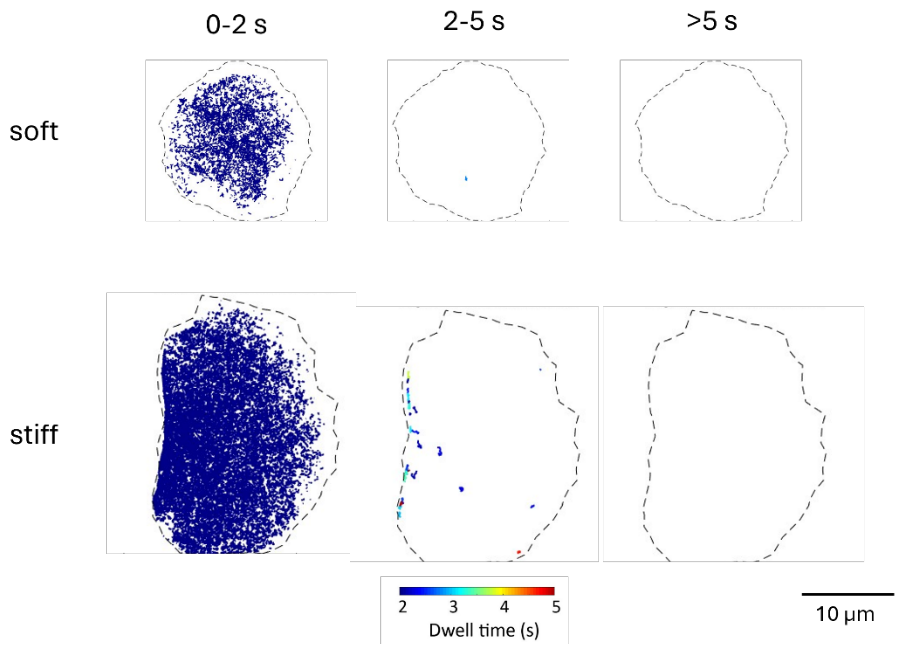


Figure S10. Representative spatial mapping of the mean mEos3.2-Grb2 dwells on soft vs stiff substrates in the starved condition. Dwells are color-coded based on duration and plotted over short (0-2s), moderate (2-5s), and long-dwelling (>5s) regimes. Scale bar, 10 µm.

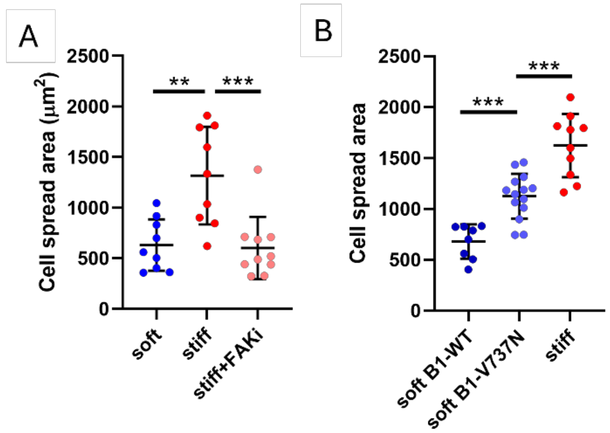


Figure S11. Cell spreading for MEC cells spread on soft vs stiff substrates with adhesion manipulations. A) Cell spreading area quantification for MEC cells spread on soft and stiff substrates with or without the FAK inhibitor (FAKi). Data are plotted with cells as scatter points with mean±s.d. Statistical significance was measured by ANOVA followed by the Kruskal-Wallis test; $**p<0.01$, $***p < 0.001$. (B Cell spreading area quantification for MEC cells spread on soft and stiff substrates with either integrin β1-WT or integrin β1-V737N mutant. Data are plotted with cells as scatter points with mean±s.d. Statistical significance was measured by ANOVA followed by the Kruskal-Wallis test; $***p < 0.001$.

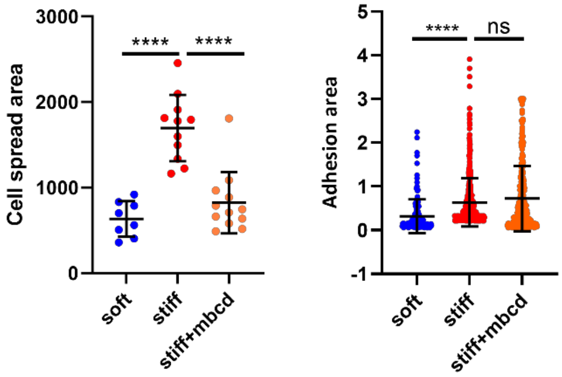


Figure S12. Cell spreading and adhesion area quantification for MEC cells spread on soft vs stiff substrates with MβCD treatment. Data are plotted with cells as scatter points with mean±s.d. Statistical significance was measured by ANOVA followed by the Kruskal-Wallis test; ns p>0.05, ****p<0.0001.

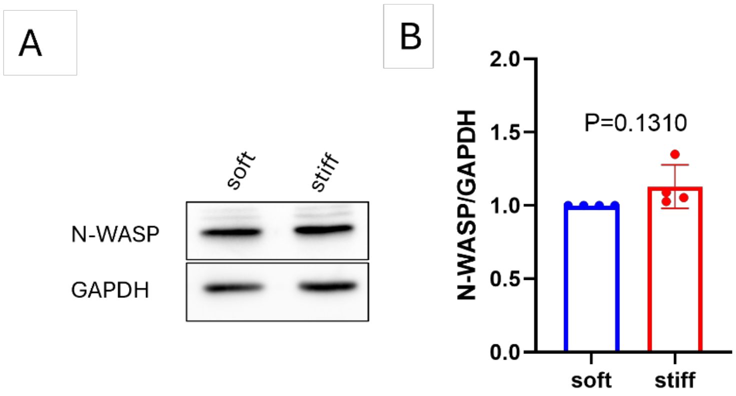


Figure S13. Substrate stiffness does not affect total N-WASP levels. A) Western blot analysis of total N-WASP levels on soft vs stiff substrates. B) Quantification of N-WASP normalized to GAPDH levels for N=4.
